# The assisting role of Polη in transcription facilitates formation of damage-induced cohesion

**DOI:** 10.1101/2020.12.20.423707

**Authors:** Pei-Shang Wu, Donald P. Cameron, Jan Grosser, Laura Baranello, Lena Ström

**Affiliations:** Karolinska Institutet, Department of Cell and Molecular Biology, SE-171 77 Stockholm, Sweden

## Abstract

The structural maintenance of chromosome (SMC) complex cohesin mediates sister chromatid cohesion established during replication, and damage-induced cohesion formed in response to DSBs post-replication. The translesion synthesis polymerase Polη is required for damage-induced cohesion through a hitherto unknown mechanism. Since Polη is functionally associated with transcription, and transcription triggers *de novo* cohesion in *Schizosaccharomyces pombe*, we hypothesized that transcription facilitates damage-induced cohesion in *Saccharomyces cerevisiae*. Here, we show dysregulated transcriptional profiles in Polη-depleted cells (*rad30Δ*), where genes involved in chromatin assembly and positive transcription regulation were downregulated. In addition, chromatin association of RNA polymerase II was reduced at promoters and coding regions in *rad30Δ* compared to WT cells, while occupancy of the H2A.Z variant (Htz1) at promoters was increased in *rad30Δ* cells. Perturbing histone exchange at promoters inactivated damage-induced cohesion, similarly to deletion of the *RAD30* gene. Conversely, altering regulation of transcription elongation suppressed the deficient damage-induced cohesion in *rad30Δ* cells. These results indicate that Polη has an assisting role during the transcriptional process, which consecutively facilitates formation of damage-induced cohesion. This further suggests a potential linkage between regulation of transcription and formation of damage-induced cohesion after replication.

**Author Summary:** The cohesin complex dynamically associates with chromosomes and holds sister chromatids together through cohesion established during replication. This ensures faithful chromosome segregation at anaphase. In budding yeast, DNA double strand breaks trigger sister chromatid cohesion even after replication. This so-called damage-induced cohesion is formed both close to the breaks, and genome-wide on undamaged chromosomes. The translesion synthesis polymerase eta (Polη) is specifically required for genome wide damage-induced cohesion. Although Polη is well characterized for its function in bypassing ultraviolet-induced DNA lesions, its mechanistic role in damage-induced cohesion is unclear. Here, we show that transcriptional regulation is perturbed in the absence of Polη. We propose that Polη could aid in chromatin association of RNA polymerase II through phosphorylation of the Polη-S14 residue, a non-canonical role of Polη which further facilitates formation of damage-induced cohesion genome wide. In addition, we observe the need of replication-independent nucleosome assembly/histone exchange for formation of damage-induced cohesion. This together provides new insight into formation of damage-induced cohesion after replication, which will be interesting to further explore.

## Introduction

Dynamic disassembly and reassembly of nucleosomes — the building blocks of chromatin — facilitates processes such as replication and transcription. During the course of chromatin assembly, the canonical histones are exchanged with histone variants or post-translationally modified histones. This affects the physical and chemical properties of nucleosomes, as well as chromatin accessibility. Replication-independent nucleosome assembly, or so-called histone exchange, aids and regulates RNA polymerase II (RNAPII) passage through the nucleosomes during transcription initiation and elongation [1]. This is accomplished through histone chaperones, in concert with histone modifying enzymes and chromatin remodelers [2].

Transcription is not only the instrument for gene expression, but is also connected to cohesin localization on chromosomes. Cohesin is one of the structural maintenance of chromosomes (SMC) protein complexes, with the core formed by Smc1, Smc3 and the kleisin Scc1. Cohesin dynamically associates with chromosomes at intergenic regions of convergent genes, possibly as a result of active transcription [3, 4]. Cohesin and its chromatin loader Scc2 have been implicated in gene regulation [5–7] and also in spatial organization of chromosomes into topologically associated domains (TADs) through DNA loop extrusion [8–12].

In addition to the roles described above, the canonical role of cohesin is to mediate sister chromatid cohesion. Cohesin is recruited to chromatin by the cohesin loading complex Scc2-Scc4 from late G_1_ phase in *S. cerevisiae* [13], and continuously through the cell cycle [14, 15]. During S-phase, cohesin becomes cohesive through acetylation of Smc3 by the acetyltransferase Eco1 [16–18]. The established sister chromatid cohesion is then maintained until anaphase [19], ensuring faithful chromosome segregation.

At the end of S phase, Eco1 is targeted for degradation. However, inducing a single site-specific double strand break post-replication (G_2_/M) is sufficient to stabilize Eco1 [20, 21]. Presence of active Eco1 then allows generation of damage-induced cohesion in G_2_/M, which is established close to the break, and also genome wide on undamaged chromosomes [22–24]. We previously showed that Polymerase eta (Polη), one of the three translesion synthesis (TLS) polymerases in *S. cerevisiae*, is specifically required for genome wide damage-induced cohesion [25].

Polη (encoded by the *RAD30* gene) is well characterized for bypassing bulky lesions induced by ultraviolet irradiation [26], yet emerging evidence suggest that Polη also exhibits TLS-independent functions [27]. Polη is the only TLS polymerase required for damage-induced cohesion [25], independently of its polymerase activity, but dependent on Polη-S14 phosphorylation; potentially mediated by the cyclin dependent kinase, Cdc28 [28]. However, the underlying role of Polη in damage-induced cohesion remains unclear. Thus, absence of Polη does not affect break-proximal damage-induced cohesion or DSB repair. Lack of Polη also does not perturb Eco1 stabilization, cohesin chromatin association or Smc3 acetylation after induction of DSBs in G_2_/M [25].

Based on the following two observations, we hypothesized that active transcription facilitates damage-induced cohesion genome wide. First, Polη is enriched at actively transcribed regions, and required for expression of several active genes in *S. cerevisiae* [29]. Second, activated transcription leads to establishment of local *de novo* cohesion in *S. pombe* [30]. Here, we present data pointing at a role for Polη and Polη-S14 phosphorylation in facilitating chromatin association of RNAPII. In addition, the transcriptional program in the Polη null mutant (*rad30Δ*) is altered both before and after DSB induction, with expression of genes involved in chromatin assembly and positive transcription regulation being downregulated compared to WT cells. Perturbing histone exchange at promoter regions by a *HIR1* or *HTZ1* deletion negatively affects damage-induced cohesion formation, in a similar fashion as in *rad30Δ* cells. Deletion of the transcription elongation regulator *SET2* however, suppresses the lack of damage-induced cohesion in the *rad30Δ* mutant. Taken together, our results suggest that Polη is required for damage-induced cohesion through its assisting role in transcription, and support the hypothesis that regulated transcription facilitates formation of damage-induced cohesion.

## Results

### A potential role for Polη in facilitating chromatin association of RNAPII

To test if active transcription is correlated with generation of damage-induced cohesion, we initially assessed sensitivity of the damage-induced cohesion deficient *rad30Δ* and *Polη-S14A* cells to transcription elongation inhibitors. Viability of both mutants decreased when exposed to actinomycin D (Fig 1A). In addition, consistent with a previous report [29], *rad30Δ* cells were sensitive to mycophenolic acid (MPA). This was also true for the *Polη-S14A* point mutant (Fig 1A). Sensitivity of both mutants to MPA was reversed by supplementing guanine in the media (Fig 1A), verifying that it was due to depletion of the guanylic nucleotide pool [31].

**Fig 1.**
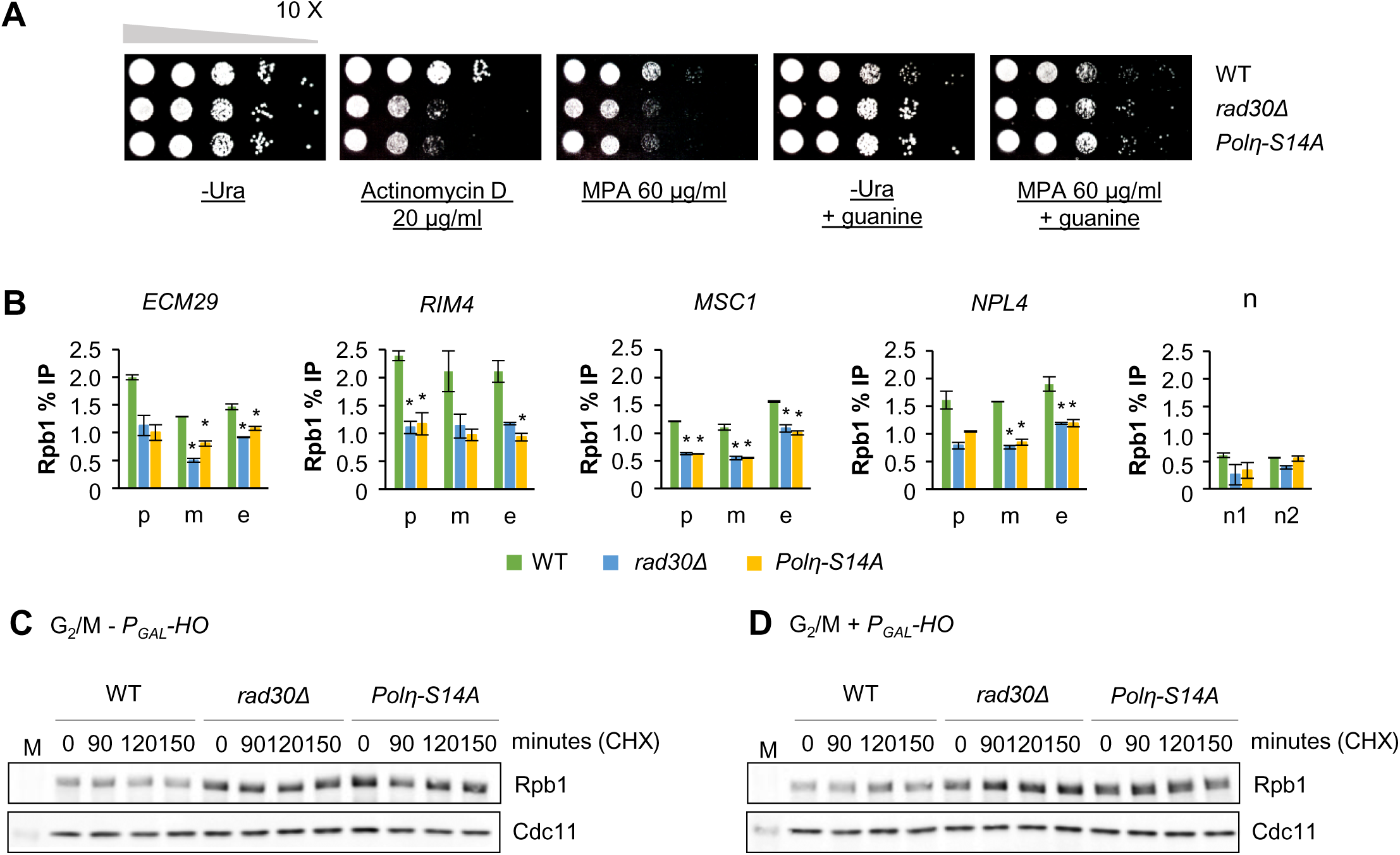
A potential role for Polη in facilitating chromatin association of RNAPII. (A) Spot assay to monitor sensitivity of the *rad30Δ* and *Polη-S14A* mutants to the transcription elongation inhibitors, actinomycin D and mycophenolic acid (MPA). Tenfold serial dilutions of indicated mid-log phase cells on controls (-Ura plate ± guanine), and drug-containing plates, after 3 days incubation at room temperature. (B) ChIP-qPCR analyses to determine chromatin association of Rpb1 in indicated strains, on selected actively transcribed genes in G_2_/M arrested WT cells. Error bars indicate the mean ± STDEV of two independent experiments. Asterisks denote significant differences compared to the WT cells at indicated position (*p* < 0.05; One-way ANOVA, Tukey post hoc test). p, promoter; m, mid; e, end of gene body. n1 and n2, low-binding controls. (C-D) Western blot analysis of Rpb1 stability. G_2_/M arrested cells from indicated strains, with or without one-hour P*_GAL_-HO* break induction, were pelleted and resuspended in media containing cycloheximide (CHX) to monitor Rpb1 protein levels without further protein synthesis. Cdc11 was used as loading control. M, protein marker.

Sensitivity to elongation inhibitors might be due to reduced transcriptional capacity. We therefore monitored chromatin association of Rpb1, the largest subunit of RNAPII, in these mutants. Binding of Rpb1 at promoters and coding regions of selected active genes was reduced in both *rad30Δ* and *Polη-S14A* mutants compared to WT cells (Fig 1B). The reduced chromatin association was accompanied by an increased level of total Rpb1 (Figs 1C and S1A). Furthermore, Rpb1 stability in the *rad30Δ* and *Polη-S14A* mutants was not affected, regardless of DSB induction (Figs 1C and 1D, S1A and S1B). Here and throughout the study the DSBs were induced at the *MAT* locus on chromosome III (P*_GAL_-HO*) for one-hour, unless otherwise stated. These results together suggest that Polη facilitates chromatin association of RNAPII for proper transcription initiation and elongation, independent of DNA damage, likely through phosphorylation of Polη-S14.

### Transcription is perturbed in *rad30Δ* mutants

To further pinpoint a potential connection between transcription and formation of damage-induced cohesion, we focused on the *rad30Δ* mutant for the following investigations. To begin with, we analyzed gene expression of G_2_/M arrested WT and *rad30Δ* cells, before and after one-hour break induction, by RNA-sequencing analysis (RNA-seq). Prior to RNA-seq, G_2_/M arrest and break induction were confirmed (S2A and S2B Fig). Principal component analysis (PCA) showed that the individual data sets were distributed as distinct clusters (S2C Fig). Differences in gene expression patterns between WT and *rad30Δ* cells were readily observed before break induction, with 395 genes upregulated and 439 genes downregulated in the G_2_/M arrested *rad30Δ* mutant (Fig 2A). In response to DSB induction, the WT cells showed 473 genes up- and 519 genes down-regulated (Fig 2B), whereas there were 360 genes up- and 230 genes down-regulated in the *rad30Δ* mutant (Fig 2C, S1 Data). While the differentially expressed genes in WT and *rad30Δ* cells after break induction significantly overlapped (S2D Fig) and trended in the same direction, the up- and down-regulation after DSB was of greater magnitude in the WT cells (Fig 2D and 2E). This implied that the response to break induction in the *rad30Δ* cells is similar, but relatively attenuated in comparison to the response in WT cells. Furthermore, we noted that short genes were preferentially upregulated compared to long genes in WT cells after DSB induction (Fig 2F), similar to the reported gene length dependent changes of expression after UV exposure [32, 33]. In contrast, differential expression after DSBs is independent of gene length in the *rad30Δ* mutant (Fig 2F). From these results we conclude that *RAD30* deletion leads to transcription deregulation, both in unperturbed G_2_/M phase and in response to break induction.

**Fig 2.**
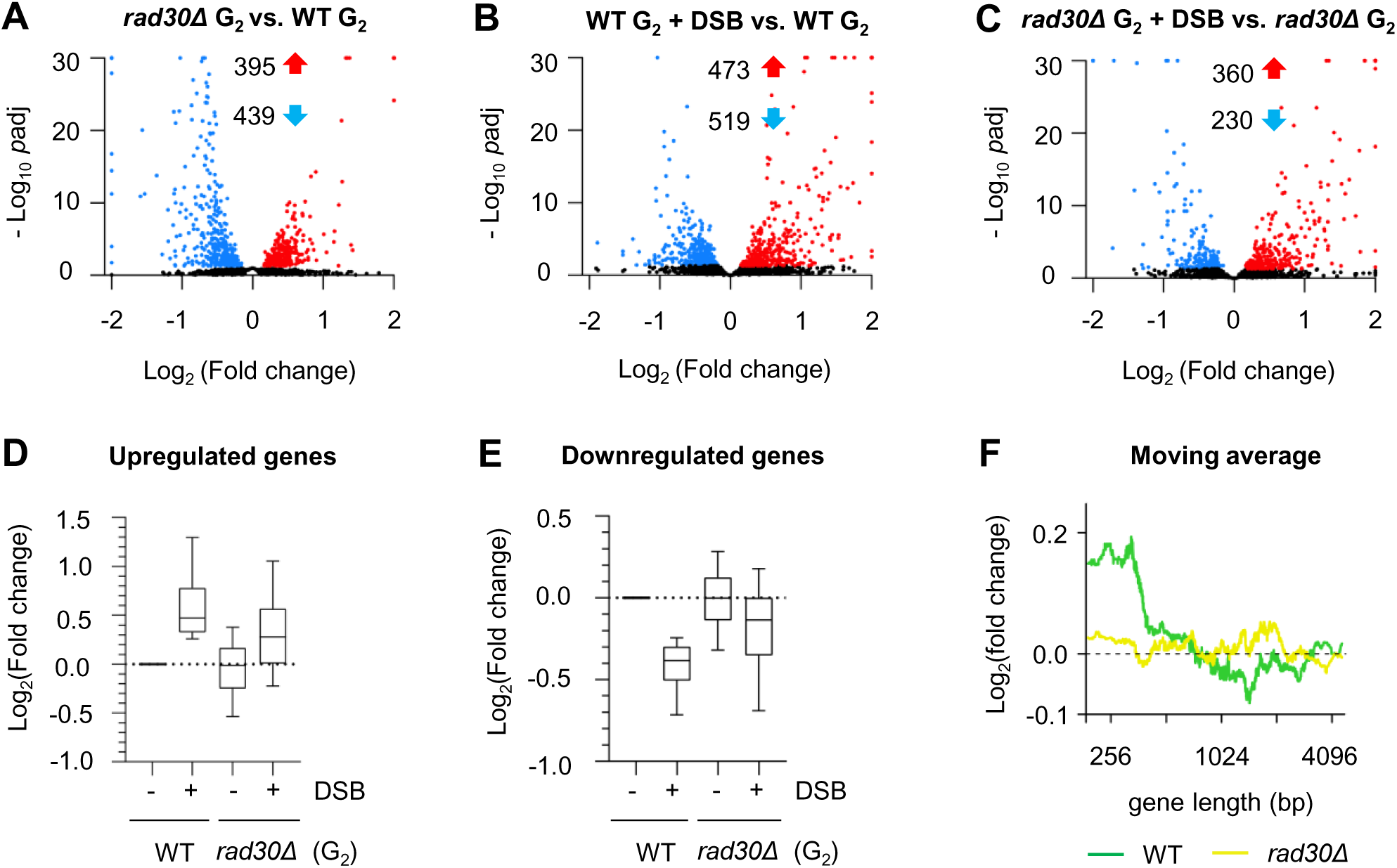
Transcription is perturbed in *rad30Δ* mutants. (A-C) Volcano plots showing differentially expressed genes between WT and *rad30Δ* cells, before and after DSB, determined by RNA-seq. Each dot represents one gene. Red and blue dots represent up- and down-regulated genes respectively. Numbers of differentially expressed genes are indicated. Black dots indicate genes without significant changes in expression (*p*adj < −Log_10_(0.5)). *p*adj, adjusted *p* value. (D-E) Comparisons between expression level of genes significantly up (D) or downregulated (E) in the WT+DSB relative to the G_2_/M arrested WT cells, and expression of the same set of genes in the *rad30Δ* mutant, based on RNA-seq analysis. (F) Plot of fold change moving median, sorted by length (300 genes/window) to monitor the trend of gene expression after DSB in relation to gene length, comparing WT and *rad30Δ* cells. Fold change values were based on the changes of gene expression in WT and *rad30Δ* cells after DSB, determined by RNA-seq.

### Downregulated genes in G_2_/M arrested *rad30Δ* cells are enriched for closed- and TATA-containing promoters

To gain additional insight into the role of Polη during transcription, we used published datasets to analyze if the deregulated genes in *rad30Δ* cells were associated with specific types of promoters, in a similar manner as reported [34]. These datasets classify genes according to type of promoter: (i) open/closed promoters, either with or without a nucleosome free region [35], (ii) promoters with fragile/stable nucleosome, defined by sensitivity of the −1 nucleosome to MNase digestion [36], and (iii) the canonical TATA-containing or TFIID dominated promoters [37, 38]. Notably, a significant number of downregulated genes in G_2_/M arrested *rad30Δ* cells were classified under the group of closed promoters (Table 1). In addition, the up- and down-regulated genes in G_2_/M arrested *rad30Δ* cells were dominated by TATA-containing promoters (obs/exp>1). These imply that Polη more frequently associates with promoters in closed configuration and TATA-containing promoters, primed for transcriptional activation in G_2_/M phase.

**Table 1.** Association of differentially expressed genes with promoter type in G_2_/M arrested *rad30Δ* cells

| *rad30Δ* G2 vs. WT G2 | upregulated (395) |  |  | downregulated (439) |  |  |
| --- | --- | --- | --- | --- | --- | --- |
|  | overlap | obs/exp | p values | overlap | obs/exp | p values |
| closed promoter (1596) | 118 | 1.1 | 0.046 | 146 | **1.3** | 3.309e-04 * |
| open promoter (3504) | 228 | 1.0 | 0.459 | 237 | 0.9 | 0.077 |
| FN promoter (1953)<sup>a</sup> | 139 | 1.1 | 0.086 | 156 | 1.1 | 0.054 |
| SN promoter (3066)<sup>b</sup> | 206 | 1.0 | 0.223 | 245 | 1.1 | 0.008 |
| TATA-containing (1090) | 96 | **1.4** | 6.726e-04 * | 132 | **1.7** | 5.069e-11 * |
| TFIID-dominated (5130) | 299 | 0.9 | 7.636e-06 * | 326 | 0.9 | 4.377e-08 * |

Number of genes in each group is indicated in parentheses. The numbers in bold indicate that the overlap is higher than expected, observation/expectation (obs/exp)>1. Asterisks indicate significant overlap ( $p<0.001$ ), evaluated as described in materials and methods. <sup>a</sup>FN: fragile nucleosome, <sup>b</sup>SN: stable nucleosome.

### Increased cohesin binding around TSS in *rad30Δ* cells

Since active transcription results in cohesin localization at the ends of convergent genes [3, 39], we considered that the observed transcriptional deregulation in the *rad30Δ* mutant could affect cohesin dynamics on chromosomes. To address this possibility, we re-analyzed our previously published Scc1 ChIP-sequencing dataset (GSE42655), from which it was concluded that cohesin binding was similar with and without break induction in the WT cells, except at the break site. In addition, absence of Polη did not result in apparent differences in overall cohesin binding [25]. The upregulation of short genes in WT cells after DSB induction (Fig 2F) also appeared to be independent of cohesin binding (S2E Fig). Thus, transcription responses in general do not seem to directly correlate with cohesin distribution in G_2_/M phase. However, when focusing on cohesin binding at transcription start site (TSS) and transcription end site (TES), accumulation of cohesin around TSS was increased in *rad30Δ* compared to WT cells (Fig 3A and 3B). Notably, this increased accumulation was not found around TES (Fig 3C and 3D), and was independent of DSB induction (Fig 3A-D). This could reflect that cohesin associated around TSS becomes less dynamic when transcription is dysregulated, regardless of DSB induction.

**Fig 3.**
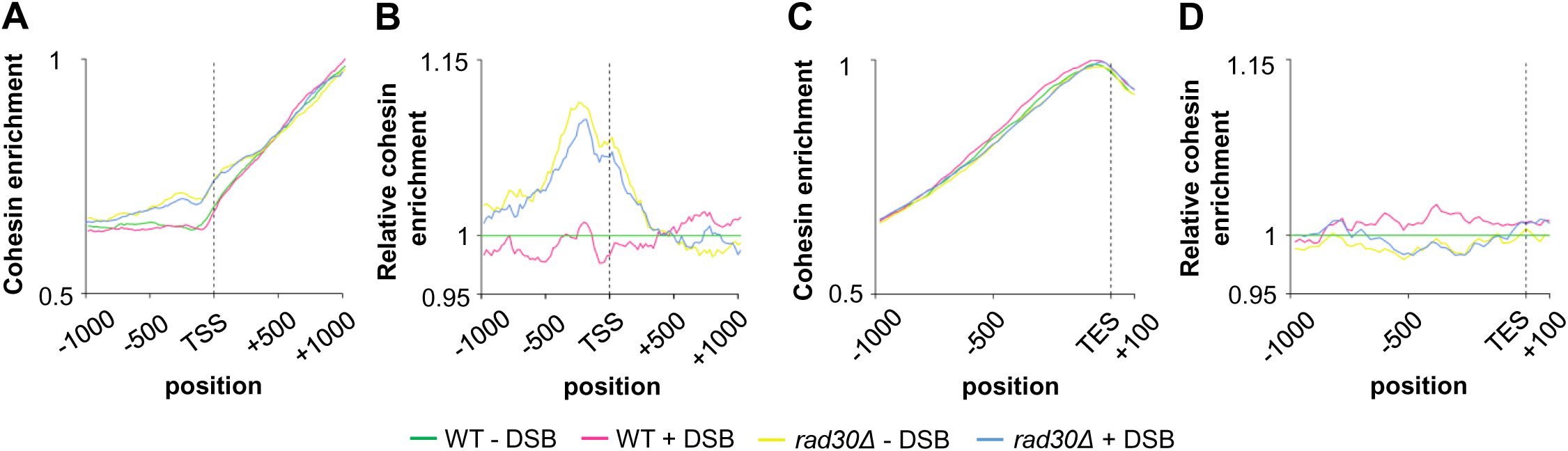
Increased cohesin binding around TSS in *rad30Δ* cells. (A) Metagenome plot showing cohesin enrichment ± 1000 bp from the transcription start site (TSS) in WT and *rad30Δ* cells ± DSB induction in G_2_/M phase. The samples were first normalized to their respective input and then the values were scaled to the maximum value of the plot. (B) The data from (A) plotted relative to the WT-DSB sample. After normalizing to the input, all samples were also normalized to WT-DSB sample to visualize the changes between the WT and *rad30Δ* cells. (C) Metagenome plot showing cohesin distribution 1000 bp downstream and 100 bp upstream from the transcription end site (TES) in WT and *rad30Δ* cells ± DSB induction in G_2_/M phase. Plotted as in (A). (D) As in (B), except plotting cohesin distribution around the TES according to (C).

### Genes involved in chromatin assembly and positive transcription regulation pathways are downregulated in the absence of Polη

To gain mechanistic insight into the diverse transcriptional responses detected in WT and *rad30Δ* cells, differential gene expression between WT and *rad30Δ* cells (before and after DSBs) were analyzed by Gene Set Enrichment Analysis (GSEA), followed by generation of enriched pathway maps with Cytoscape as shown in Fig 4. The gene sets under each annotated group are listed in S2 and S3 Data. During G_2_/M arrest, genes that belong to biological pathways such as chromatin assembly and positive transcription regulation were downregulated in *rad30Δ* compared to WT cells (Fig 4A). Consistent with downregulation of the genes involved in chromatin assembly pathway, we observed that the nucleosome occupancy of *rad30Δ* cells was moderately increased compared to WT cells (S3A Fig). When comparing gene expression after break induction, the pathways illustrated in Fig 4B were clearly differentially regulated between WT and *rad30Δ* cells. WT cells tended to downregulate essential cell homeostatis pathways, such as ribosome biogenesis and various metabolism pathways, relative to the *rad30Δ* mutant. This further indicates deregulation of gene expression in the *rad30Δ* mutant. Despite that some genes belonging to the cellular response to DNA damage stimulus pathway (GO: 6974) were upregulated in WT cells after DSB induction, this pathway was overall not significantly enriched. In addition, activation of the DNA damage checkpoint, as indicated by phosphorylation of Rad53, was only observed during the recovery period after DSB induction in WT and *rad30Δ* cells (S3B and S3C Fig), with no difference in cell cycle progression between populations (S3D Fig). These results indicate that the lack of damage-induced cohesion in *rad30Δ* cells is not due to a possible difference in activation of the DNA damage checkpoint. Furthermore, in response to DSBs, expression of the acetyltransferase *ECO1* was not enhanced in either WT or *rad30Δ* cells (S3E Fig). Based on this, it is plausible to test the potential connection between transcription and damage-induced cohesion, by focusing on two of the upregulated gene sets in the WT cells before DSB induction — chromatin assembly and positive transcription regulation.

**Fig 4.**
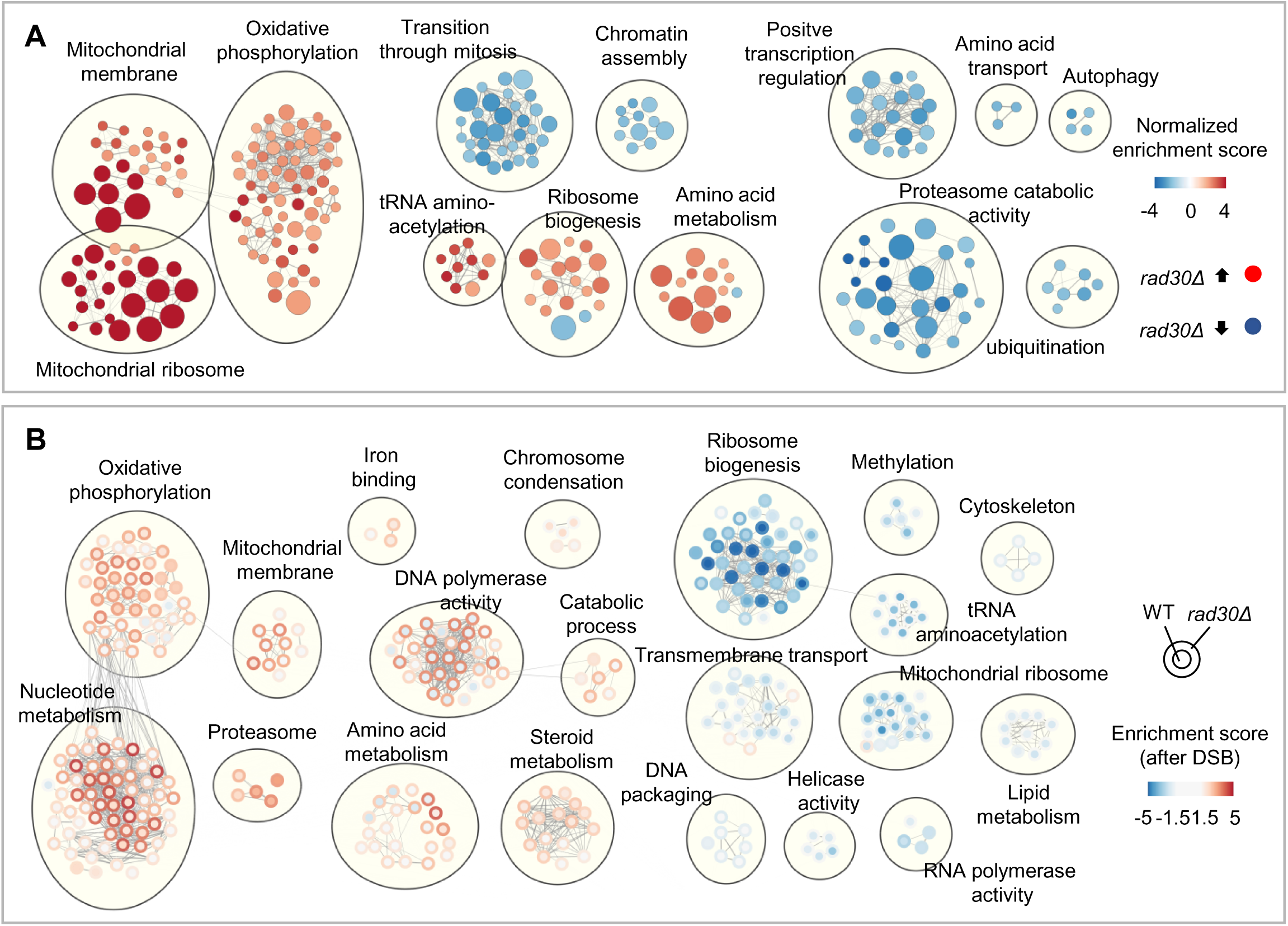
Genes involved in chromatin assembly and positive transcription regulation pathways are downregulated in the absence of Polη. (A) Relatively enriched pathways in G_2_/M arrested WT and *rad30Δ* cells, plotted with Cytoscape after gene set enrichment analysis (GSEA). The GSEA was performed with gene lists ranked by log_10_ *p* value (multiplied by the sign of the fold change) of each gene. The number of genes in each gene set is proportional to the circle size. Lines connect gene sets with similarity greater than 0.7. All gene sets have FDR < 0.05. (B) Gene set enrichment analysis after DSB induction, plotted with Cytoscape to depict the difference between WT and *rad30Δ* cells in up- or down-regulation of indicated pathways after DSB. Gene expression of WT and *rad30Δ* cells after DSB was compared to that of respective G_2_/M arrested cells. GSEA was performed as in (A). The lines indicate the same as in (A). All gene sets have FDR < 0.05 and a normalized enrichment score > 2 for at least one of the WT or *rad30Δ* cells.

### Deleting *HIR1* leads to partially deficient damage-induced cohesion

To test if active transcription facilitates formation of damage-induced cohesion genome-wide, we treated cells briefly with the transcription inhibitor thiolutin to block transcription before γ-irradiation. However, we observed that the thiolutin treatment itself triggered an early DNA damage response (S4A Fig). Since the γ-H2AX signal was comparable to that in irradiated cells without thiolutin (S4A Fig), we concluded that using thiolutin in our damage-induced cohesion assay is not applicable. Therefore, to test if transcriptional activity is related to generation of damage-induced cohesion, we set out to utilize a genetic approach by testing mutants which would either mimic or reverse the transcriptional deregulation in *rad30Δ* cells.

To this end, from the perspective of chromatin assembly, we investigated whether Hir1 (a component of the HIR complex) is required for damage-induced cohesion. The HIR complex and the histone chaperone Asf1 mediate histone H3 exchange with post-translationally modified H3, independently of replication [40, 41]. The exchange mainly takes place at promoters and correlates with active transcription. However, basal H3 exchange also occurs to poise inactive promoters for optimal transcription [42, 43]. To monitor damage-induced cohesion, DSBs and ectopic P*_GAL_-SMC1-MYC* expression were induced by addition of galactose to G_2_/M arrested cells. Due to the *smc1-259 ts* background, cohesion established during replication was inactivated by raising the temperature. Damage-induced cohesion generated with the ectopic Smc1-Myc was examined with an integrated TetO/TetR-GFP array on Chr. V (illustrated in S4B Fig). G_2_/M arrest, break induction and protein expression of the ectopic Smc1-Myc were confirmed for all experiments, with examples shown in S4C-S4E Fig. Interestingly, formation of damage-induced cohesion was partially deficient in the *hir1Δ* mutant, while the *hir1Δrad30Δ* double resembled the *rad30Δ* single mutant, although with slower sister separation (Fig 5A). This indicated that Hir1 and Polη are both required for efficient damage-induced cohesion; possibly acting in the same pathway.

**Fig 5.**
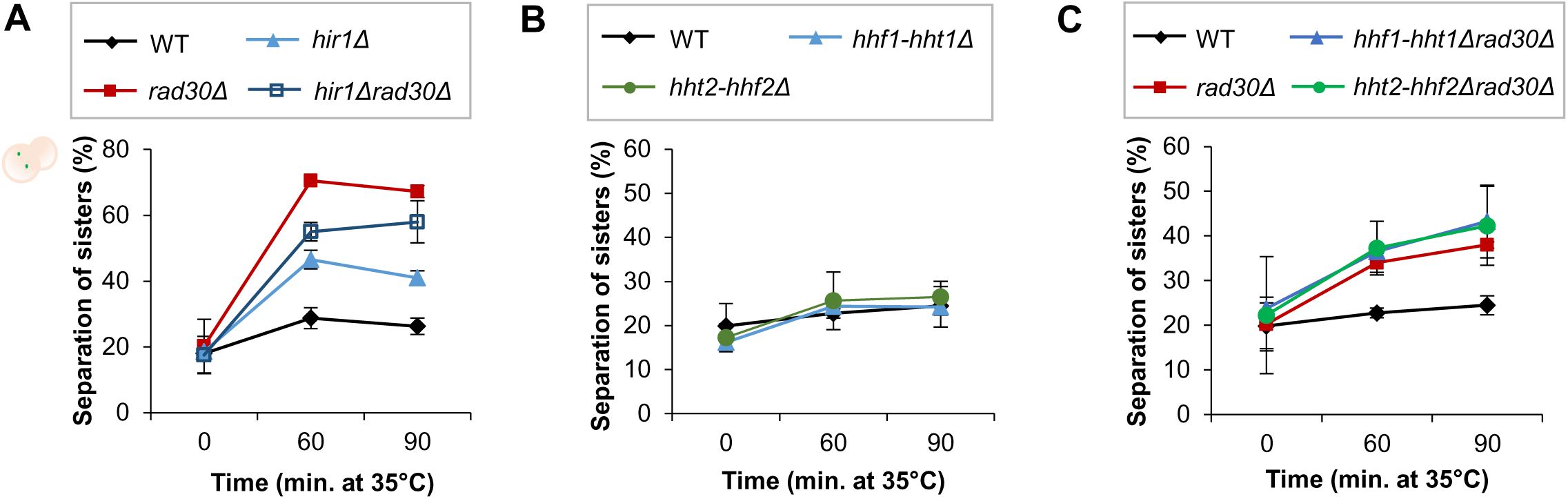
Deleting *HIR1* leads to partially deficient damage-induced cohesion. (A) Damage-induced cohesion assays of the *hir1Δ* single and *hir1Δrad30Δ* double mutants after P*_GAL_*-*HO* induction, performed as illustrated in S4B Fig. Means ± STDEV from at least two independent experiments are shown. (B-C) Damage-induced cohesion assays of the *hhf1-hht1Δ* and *hht2-hhf2Δ* mutants after P*_GAL_*-*HO* induction, performed as in (A). Means ± STDEV from at least two independent experiments are shown.

With the *hir1Δ* mutant we aimed at testing if the role of the HIR complex in chromatin assembly affected formation of damage-induced cohesion. However, the observed deficiency of the *hir1Δ* cells might be due to de-repression of histone genes, as the HIR complex also negatively regulates histone genes expression [44, 45]. If so, reducing the histone gene dosage should be beneficial for the *rad30Δ* mutant in generation of damage-induced cohesion. Yet, deletion of any H3-H4 coding gene pair (*HHT1-HHF1* and *HHT2-HHF2*) did not affect formation of damage-induced cohesion in *rad30Δ* cells (Fig 5B and 5C). This indicates that the partial deficiency of the *hir1Δ* mutant is not due to altered histone gene dosage, and points to a need for histone exchange during transcription for formation of damage-induced cohesion.

### Perturbing histone exchange at promoters negatively affects formation of damage-induced cohesion

To further investigate the effect of pertubing histone exchange on formation of damage-induced cohesion, we tested requirement of the H2A variant Htz1 (H2A.Z) in this process. Htz1 is preferentially incorporated at basal/repressed promoters. Susceptibility of Htz1 to loss from the incorporated nucleosome promotes its exchange for H2A. This facilitates transcriptional activation [46, 47], and relieves the +1 nucleosome barrier to RNAPII [48, 49]. Since the *htz1Δ* mutant does not respond to P*_GAL_-HO* induction [50], γ-irradiation was utilized as source of DSB induction (see materials and methods). Similar to the *hir1Δ* mutant (S5A and S5B Fig), the *htz1Δ* mutant showed impaired damage-induced cohesion (Fig 6A). We noted that in contrast to a previous report [51], we did not observe a cohesion maintenance defect due to *HTZ1* deletion (S5C Fig).

**Fig 6.**
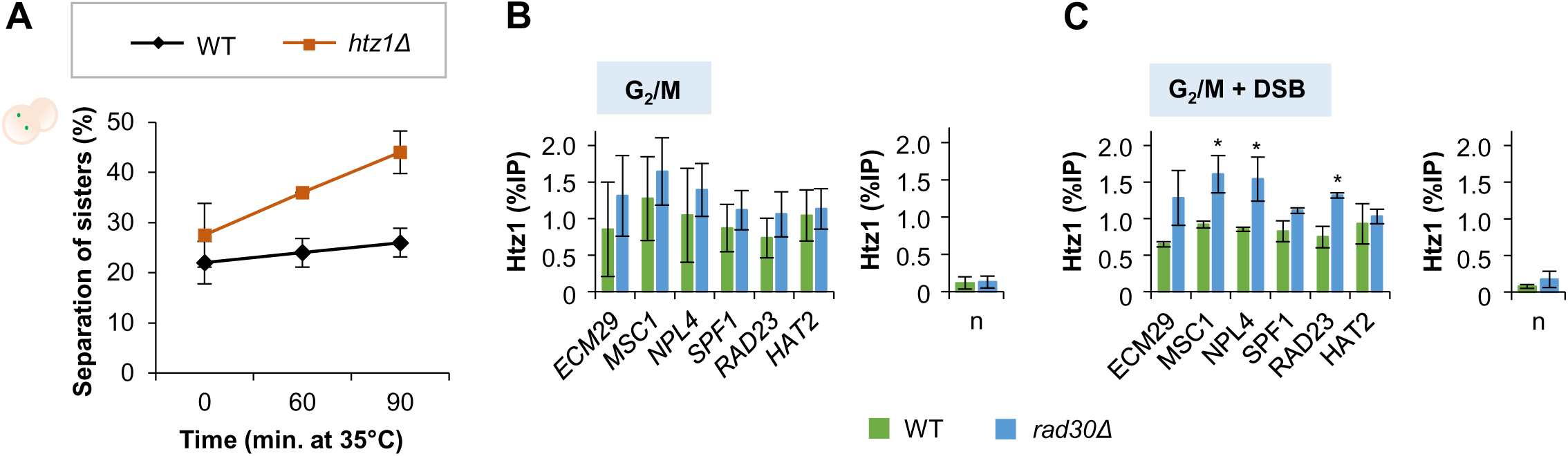
Perturbing histone exchange at promoters negatively affects formation of damage-induced cohesion. (A) Damage-induced cohesion assay of the *htz1Δ* mutant after γ-irradiation, performed according to the procedure described in the materials and methods. Means ± STDEV from at least two independent experiments are shown. (B-C) ChIP-qPCR analyses to determine Htz1 occupancy at promoters of selected genes, before (B) and (C) after DSB induction in G_2_/M arrested WT and *rad30Δ* cells. *SPF1*, *RAD23* and *HAT2* are located at the left arm of chromosome V, where damage-induced cohesion was monitored. Error bars indicate the mean ± STDEV of at least two independent experiments. Asterisks denote significant differences compared to the WT cells (*p* < 0.05; t-Test). n, low-binding control.

Since Htz1 is required for formation of damage-induced cohesion, we investigated if there was a difference between WT and *rad30Δ* cells in Htz1 occupancy at promoters. For this, we focused on the active genes analyzed for Rpb1 binding in Fig 1B and a few genes around the *URA3* on Chr. V, where we monitored damage-induced cohesion. We further selected genes with TATA-less promoters for analyses because Htz1 is relatively enriched at these promoters [46, 47]. Interestingly, Htz1 occupancy at some of the selected promoters was significantly increased in *rad30Δ* compared to WT cells, particularly after DSB induction in G_2_/M (Fig 6B and 6C). This indicates that the Htz1/H2A exchange at certain promoters was reduced in the absence of Polη, especially in response to DSB. These results were in line with *hir1Δ* and *htz1Δ* cells being deficient in damage-induced cohesion (Figs 5A, 6A and S5B), and suggest that perturbing histone exchange at promoters negatively affects formation of damage-induced cohesion.

### The assisting role of Polη in transcription is needed for generation of damage-induced cohesion

In addition to the *hir1Δ* and *htz1Δ* mutants, we utilized a *set2Δ* mutant to test if transcriptional regulation is correlated with generation of damage-induced cohesion. Set2 mediates co-transcriptional H3K36 methylation (H3K36me2/3). This promotes restoration of chromatin to the pretranscribed hypoacetylation state and represses histone exchange at coding regions during transcription elongation [52–54]. Presence of Set2 at promoters also suppresses transcription initiation of certain basal repressed genes [55–57]. Interestingly, a *set2Δ* mutant suppressed sensitivity of certain transcriptional elongation factor mutants to 6-azauracil [57], a mechanistic analog of MPA [58, 59]. As we showed that *rad30 Δ* cells are sensitive to transcription elongation inhibitors (Fig 1A), we tested if deletion of *SET2* would rescue *rad30Δ* cells from being sensitive to these inhibitors. The *set2Δ* mutant showed no obvious sensitivity to MPA or actinomycin D, and masked the sensitivity of *rad30Δ* cells especially to actinomycin D (Fig 7A). This suggests that Set2 could counteract Polη during transcription elongation. Through this genetic interaction, we tested if deletion of *SET2* would also suppress the deficiency of *rad30Δ* cells in damage-induced cohesion. The *set2Δ* mutant resembled the WT cells in formation of damage-induced cohesion. Remarkably, deletion of *SET2* suppressed the lack of damage-induced cohesion in the *rad30Δ* mutant (Fig 7B). Given that removing *SET2* caused an increased RNAPII association towards the 3’-end of actively transcribed genes [60], we monitored chromatin association of Rpb1 in the *set2Δrad30Δ* mutant. Absence of Set2 in G_2_/M arrested *rad30Δ* cells to some extend compensated for the reduced Rpb1 binding in *rad30Δ* cells (Fig 7C-7E). This trend was however not observed after DSB induction (S6A-C Fig). Considering that the differentially expressed genes in WT and *rad30Δ* cells after DSB significantly overlapped (S2D Fig), the data together suggest that general transcriptional regulation during G_2_/M phase influences formation of damage-induced cohesion, and indicate that Polη is required for damage-induced cohesion through facilitating transcription.

**Fig 7.**
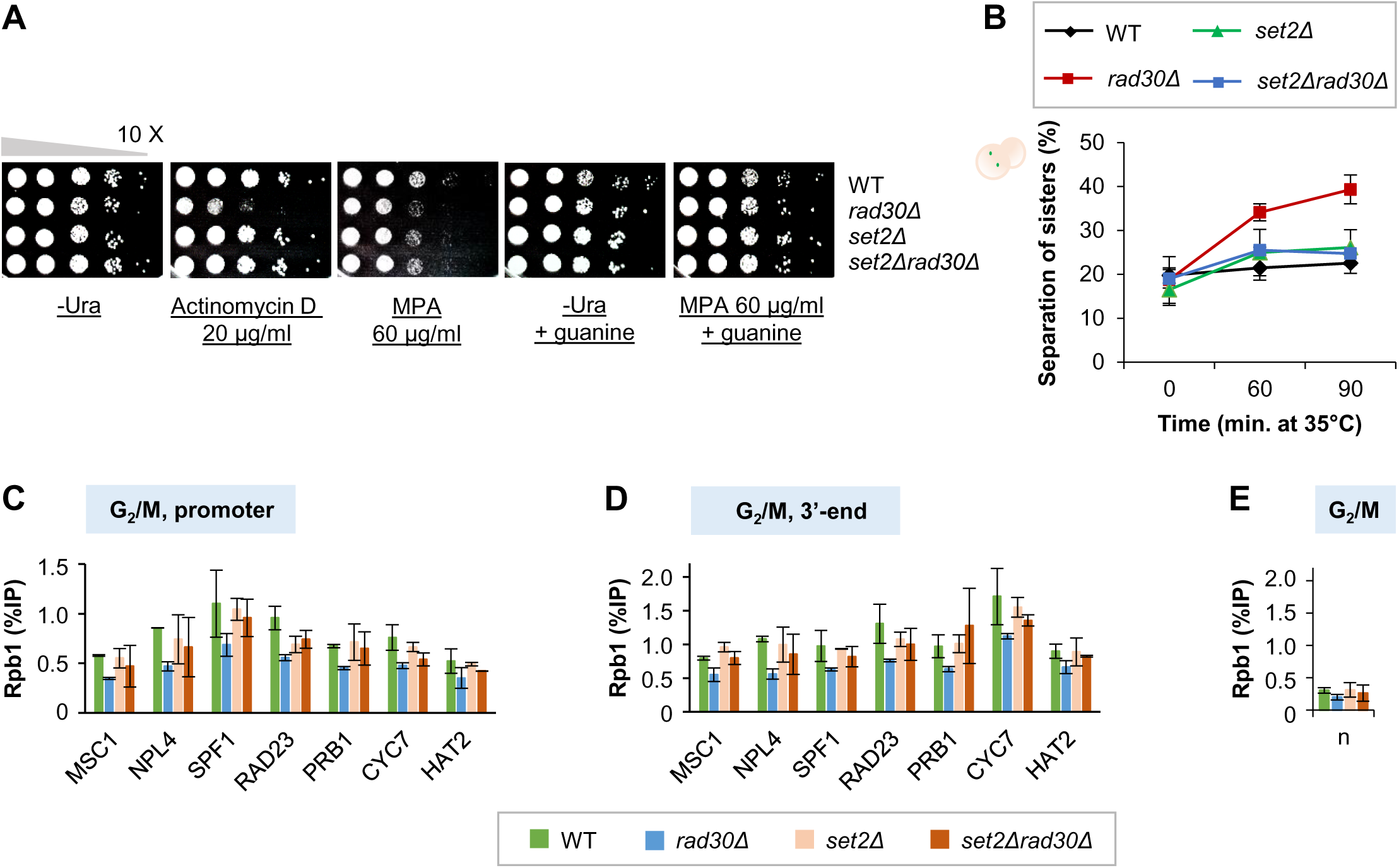
The assisting role of Polη in transcription is needed for generation of damage-induced cohesion. (A) Spot assay to monitor the effect of *SET2* deletion on the *rad30Δ* mutant sensitivity to the transcription elongation inhibitors, actinomycin D and mycophenolic acid (MPA). Tenfold serial dilutions of indicated mid-log phase cells on control (-Ura plate ± guanine) and drug-containing plates, after 3 days incubation. (B) Damage-induced cohesion assay of the *set2Δ* mutant after P*_GAL_*-*HO* induction, performed as depicted in S4B Fig. Means ± STDEV from at least two independent experiments are shown. (C-E) ChIP-qPCR analyses to determine chromatin association of Rpb1 at promoters and 3’-ends of selected genes, in indicated G_2_/M arrested cells. Except *MSC1* and *NPL4*, the rest of the selected genes are located at the left arm of chromosome V, where damage-induced cohesion was monitored. Error bars indicate the mean ± STDEV of at least two independent experiments. n, low-binding control (n2 in Fig 1B).

## Discussion

We previously showed that Polη is specifically required for genome wide damage-induced cohesion [25] but its mechanistic role in this process was unclear. This study was initiated by the observation that Polη-deficient cells displayed altered transcriptional regulation, both in unchallenged G_2_/M arrested cells and in response to DSBs. Transcription elongation deficiency was corroborated by increased sensitivity of Polη-deficient cells to transcription elongation inhibitors (Fig 1A). It could be argued that the sensitivity to actinomycin D would be a consequence of DNA damage because actinomycin D also inhibits topoisomerases [61], leading to formation of DSBs. However, since the *rad30Δ* mutant is insensitive to specific topoisomerase inhibitors, such as camptothecin and etoposide [62, 63], this was less likely.

To know which pathways were affected in the absence of Polη, gene set enrichment analysis was performed after RNA-seq. We found that mitochondrial related pathways were enhanced in *rad30Δ* cells, in contrast to downregulation of genes belonging to the chromatin assembly pathway (Fig 4A and 4B, S2 and S3 Data). This is an interesting observation since genes involved in the tricarboxylic acid cycle and oxidative phosphorylation pathways, which are related to mitochondria, were similarly upregulated in mutants with defective chromatin assembly [64].

To test the idea that the lack of damage-induced cohesion in *rad30Δ* cells would be due to transcriptional dysregulation, we began by testing the requirement of HIR/Asf1 mediated histone exchange for damage-induced cohesion, from the perspective of chromatin assembly. By deleting the *HIR1* gene, which is sufficient to disrupt the HIR/Asf1 interaction [41], we found that the *hir1Δ* mutant is partially deficient in damage-induced cohesion (Figs 5A and S5B). The role of the HIR complex in damage-induced cohesion might appear difficult to pinpoint since it is involved in multiple processes. We thus addressed the possible effect of HIR-dependent repression of histone genes [44] on formation of damage-induced cohesion. This possibility was however excluded because no effect of deleting H3-H4 gene pairs (Fig 5B and 5C) was observed in *rad30Δ* cells. The HIR complex has also been implicated in formation of a functional kinetochore [65] and heterochromatic gene silencing [66]. However, the chromatin assembly complex-1 (CAF-1) is redundant with the HIR complex in these processes. Deletion of Hir1 is thereby not likely to perturb other processes than histone exchange. We therefore suggest a direct role for HIR-dependent histone exchange in damage-induced cohesion.

Functional importance of Polη in transcription was proposed to depend on its polymerase activity [29], while its role in damage-induced cohesion was not [25]. The finding that transcription facilitates formation of damage-induced cohesion could therefore be seen as conflicting with the polymerase-independent role of Polη. However, we previously showed that the putative Polη-S14 phosphorylation is required for damage-induced cohesion, but not for cell survival after UV irradiation [28], which depends on Polη polymerase activity. In addition, the *Polη-S14A* mutant exhibits similar elongation inhibitor sensitivity and altered Rpb1 behaviour as the *rad30Δ* mutant (Fig 1A-D). This together indicates that the polymerase activity is not the sole requirement for Polη in transcription.

To gain further insight into the role of Polη in transcription, we analyzed the types of promoters that Polη might associate with (Table 1). We found that the differentially expressed genes in G_2_/M arrested *rad30Δ* cells, especially the downregulated genes, were relatively enriched for closed and TATA-containing promoters. The closed promoters that lack a nucleosome free region, are known to regulate stress related genes [67]. This is consistent with the downregulation of stress response (GO:0033554) in G_2_/M arrested *rad30Δ* cells (S2 Data, Ungrouped). In addition, the TATA-box containing genes are highly regulated and associated with stress response [37]. Despite this information, it is still unclear in which way Polη facilitates chromatin association of Rpb1 (Fig 1), and how it affects Htz1/H2A exchange at promoters (Fig 6C). We speculate that Polη might serve as a scaffold protein for Rpb1 or other transcription factors to bind on chromatin, possibly through phosphorylation of the Polη-S14 residue. Further validation of the promoters that Polη preferentially associated with might provide additional clues on potential interacting transcription factors. Nevertheless, the precise role of Polη in transcription needs to be further explored.

In addition to perturbed transcriptional regulation (Fig 2), increased cohesin binding around TSS was observed in *rad30Δ* cells, independently of DSBs (Fig 3). This indicates that transcriptional regulation might also modulate the dynamics and/or positioning of cohesin at cohesin associated regions. Since the boundaries of cohesin mediated TAD-like structures often form at promoters of active genes in yeast [68], a reciprocal interplay between transcription and formation of TAD-like structure has been suggested [69]. Although this might not be directly relevant to formation of damage-induced cohesion, it would still be interesting to investigate if TAD-like structures are altered in *rad30Δ* cells, in connection with this potential reciprocal interplay.

Through perturbing histone exchange and removing a transcription elongation regulator (depicted in S7 Fig), we show that a regulated transcriptional response connected to chromatin assembly potentially facilitates generation of damage-induced cohesion post-replication. Since establishment of sister chromatid cohesion is proposed to occur simultaneously with replication fork progression [14, 70] in concert with replication-coupled nucleosome assembly [71], we propose that replication-independent nucleosome assembly could be utilized as an alternative platform for generation of damage-induced cohesion after replication (S7 Fig, WT). Deregulated transcription in *rad30Δ* cells, which perturbs histone exchange, would in turn affect formation of damage-induced cohesion (S7 Fig, *rad30Δ*).

Despite the subtle defect in chromosome segregation observed in the *rad30Δ* mutant [25], the importance of genome wide damage-induced cohesion remains to be determined. It might be relevant to the increased chromosome mobility in response to DSBs, which presumably facilitates the search of sequence homology for recombination [72, 73]. Interestingly, the movement at the same time is constrained by sister chromatid cohesion [74]. Since unbroken chromosomes are known to be less mobile than broken chromosomes [72, 73], formation of genome-wide damage-induced cohesion might further limit the movements of undamaged chromosomes, to reduce the chance of unfavorable recombinations.

In summary, we show that Polη could be an auxiliary factor for transcription and that this role facilitates formation of damage-induced cohesion. Through a genetic approach, our study provides new insight into a potential linkage between histone exchange and generation of damage-induced cohesion post-replication. Futher studies would be needed to understand how Polη aids in transcription, how chromatin dynamics during transcription facilitate formation of genome wide damage-induced cohesion, and if damage-induced cohesion could restrict movements of undamaged chromosomes.

## Materials and methods

### Yeast strains and media

All *S. cerevisiae* yeast strains, listed in S1 Table, were W303 derivatives (*ade2-1 trp1-1 can1-100 leu2-3, 112 his3-11, 15 ura3-1 RAD5 GAL psi^+^*). To create null mutants, the gene of interest was replaced with an antibiotic resistance marker through lithium acetate based transformation. Some strains were crossed to obtain desired genotypes. Yeast extract peptone (YEP) supplemented with 40 μg/ml adenine was used as yeast media, unless otherwise stated.

### Spot assay

Cell culturing and subsequent serial dilutions were performed as described. Each dilution was sequentially spotted on uracil drop-out (-Ura) media, containing actinomycin D, MPA, or solvent only (final 1.2% ethanol in plates). Guanine was supplemented at 0.3 mM final concentration [75]. The plates were kept at room temperature and documented on the third day. Each spot assay was done at least twice.

### Protein extraction and western blotting

Whole cell extracts (WCEs) were prepared with glass bead disruption, TCA or a sodium hydroxide based method [76]. To monitor Rpb1 stability, cycloheximide (Sigma) was supplemented in media (final 100µg/ml), and the protein extracts were prepared with sodium hydroxide based method. Bolt 4-12% Bis-Tris or NuPAGE 3-8% Tris-Acetate gels (Invitrogen) were used for electrophoresis, with Bolt MOPS, Bolt MES or NuPAGE Tris-Acetate SDS running buffer (Invitrogen). Proteins were transferred to nitrocellulose membranes with the Trans-blot Turbo system (Bio-Rad) or the XCell II Blot Module (Invitrogen). Antibody information is listed in the S2 Table. Odyssey Infrared Imaging and BioRad chemiluminescence system were used for antibodies detections. Image Studio Lite software was used for quantitation of protein bands.

### Chromatin immunoprecipitation (ChIP) qPCR

ChIP was in essence performed as described with some modifications [25]. Cells were crosslinked with final 1% formaldehyde for 20 minutes at room temperature, followed by addition of final 125 mM glycine for 5 minutes. The cells were washed three times in 1X cold TBS and mechanically lysed using a 6870 freezer/mill (SPEX, CertiPrep). WCEs were subjected to chromatin shearing by sonication (Bandelin, Sonopuls) for chromatin fragments of 3-500 bp. Anti-Rpb1 and anti-Htz1 antibodies were coupled to protein A and protein G Dynabeads (Invitrogen) respectively for immunoprecipitation at 4°C, overnight. Crosslinking of eluted IP and input samples was reversed, and DNA was purified. DNA analysis was performed by real time qPCR using SYBR Green (Applied Biosystems), according to manufacturer’s guidelines on an ABI Prism 7000 sequence detection system (Applied Biosystems). The genes of interest were selected based on the RNA-seq results. Primers used are listed in S3 Table. Statistical analysis was performed with SPSS statistics software (IBM).

### Total RNA extraction

For RNA-seq, G_2_/M arrested cells (about 9 OD_600_) were harvested before and after 1-hour P*_GAL_-HO* break induction. Equal amount of samples were additionally collected at each time-point as genomic DNA (gDNA) controls. The gDNA content of each sample was determined prior to total RNA extraction. Total RNA extracts were prepared with PureLink RNA Mini Kit (Invitrogen), with some modifications of the manufacture’s guidelines. Collected cell pellets were washed once with SE mix (final 1 M sorbitol and 50 mM EDTA), and resupended with 100 μl zymolyase lysis buffer (SE mix supplemented with final 3 mg/ml 100T zymolyase (Sunrise Science) and 2.5 μl Ribolock (Invitrogen). The suspension was incubated at 30°C for 60 minutes, followed by addition of 200 μl kit-provided RNA lysing buffer, supplemented with Ribolock. The rest of the procedure was performed according to the manufacture’s guidelines. To elute total RNA from columns, the volume of RNase free water for elution was adjusted according to gDNA content of each sample. For each strain, equal volume of the total RNA extract was further purified with DNA-free Kit (Invitrogen).

### RNA-sequencing and ChIP-sequencing data analyses

Total RNA samples prepared for RNA-seq (triplicates) were subsequently handled by Novogene for mRNA enrichment, library construction (250-300 bp insert cDNA library) and RNA sequencing (Illumina HiSeq X Ten, paired-end, 10 M reads). Quality controls were included for the total RNA samples and during the procedures for RNA-sequencing.

FASTQC (https://www.bioinformatics.babraham.ac.uk/projects/fastqc/) was used for quality control of the .fastq-files for both RNA- and ChIP-seq. Adapter and poor quality read trimming was performed with cutadapt [77]. The RNA-seq data was mapped with the splice-aware aligner HISAT2 [78]. The ChIP-seq data was mapped using bowtie [79] with the colorspace option enabled. Afterwards the mapped files were sorted using samtools [80]. Both sets of sequencing data were aligned to the yeast genome version SacCer3 downloaded from UCSC genome browser. Duplicates in the mapped .bam-files were removed using MarkDuplicates (http://broadinstitute.github.io/picard) from the Picard toolset.

For the RNA-seq data set, the reads were counted per gene using featureCounts [81]. The count-files were imported into R and further analyzed using edgeR [82, 83] for FPKM calculations and DESeq2 [84] for differential expression analysis. Differential expression analysis yielded fold-changes alongside significance for genes, additionally DESeq2 was used to generate principal component analysis plots. Genes with a total read count below 10 across all samples as well as those producing NAs (not available) in any of the comparisons for fold-change calculation were excluded from the analysis. As all four conditions showed a similar within-group variability in the PCA plot, for all fold-change calculations all samples were run together as opposed to subsetting the samples of interest e.g. WT G2 + DSB vs. WT G2. This allowed for more accurate estimation of the dispersion parameter and in turn calculation of significance for the fold-changes. Also, the moving average of the fold-change was calculated by ordering the genes included in the DESeq2 dataset by length and then calculating the median of a window of 300 genes around these gene. No moving average was calculated for the 75 longest and shortest genes as they did not have an even number of genes on either site for moving average calculation.

For the ChIP-seq dataset, cohesin peaks were called using MACS2 [85]. The files generated were then imported into R, where they were annotated using the package ChIPpeakAnno [86] with gene lists downloaded using the biomaRt package [87]. The lists of genes overlapping or with their gene end closest to the peak middle with cohesin peaks were read into ngs.plot [88] for metagenome analysis. After analysis had been performed, the data were replotted using the internal R plotting.

Gene set enrichment analysis (GSEA) was performed using the Broad Institute software (http://www.broad.mit.edu/gsea) [89] using *S. cerevisiae* gene sets from the Xijin Ge lab (http://ge-lab.org/#/data) [90]. The GSEA enrichment map was created using the EnrichmentMap plugin [91] for Cytoscape [92], broadly following a published protocol [93]. Groupings were facilitated by the Cytoscape AutoAnnotate plugin [94]. In the comparison of WT vs. *rad30Δ* cells, only gene sets enriched with an adjusted *p*-value of < 0.05 were plotted. In the comparison of both WT and *rad30Δ* cells ± DSB induction, only gene sets enriched with an adjusted *p*-value of < 0.05 and a normalized enrichment score (NES) > 2 for either strain were plotted.

Statistical significance of the overlapping genes in the Venn diagrams and Table 1 were calculated using either a normal approximation or the hypergeometric probability formula. The online tool on http://nemates.org/MA/progs/overlap_stats.html was used for evaluation.

### Damage-induced cohesion assay and controls

All strains used harbor the *smc1* temperature sensitive allele (*smc1-259*). The experiments with the P*_GAL_-HO* allele for DSB induction were performed as described [28], and illustrated in S4B Fig. The assay utilizing γ-irradiation as DSB source is described in S5A Fig. Considering that the *htz1Δ* mutant is benomyl sensitive [95], the strains used in this assay contain the P*_MET_-CDC20* and *smc1-259 ts* alleles. The strains were grown in methionine drop-out media (-Met) to log phase at 23°C. To arrest cells in G_2_/M phase, expression of *CDC20* was repressed by replacing the media to YEP supplemented with Met (final 2 mM) and 0.1% glucose. Galactose (final 2%) was then added for 1.5 hours to induce expression of ectopic Smc1-Myc, driven by the *GAL* promoter. The cultures were subsequently split into half and resuspended in 1X PBS. One half for γ-irradiation (250 Gy), and another half as non-irradiated control. After 1-hour recovery in YEP media supplemented with galactose and Met, the temperature was raised to 35°C and damage-induced cohesion was monitored for 90 minutes.

Proper G_2_/M arrest, expression of the ectopic Smc1-Myc and DSBs induction in these assays were confirmed with FACS analysis, western blot, and pulsed-field gel electrophoresis (PFGE) respectively. Efficiency of γ-irradiation was analyzed with Southern blot after PFGE, with a probe for chromosome XVI, as described [96].

### MNase digestion assay

G_2_/M arrested cells were crosslinked *in vivo* with formaldehyde (final 0.5%), for 20 minutes at 23°C. To quench the reaction, glycine (final 125 mM) was added in cultures for 10 minutes. The cells were then harvested and stored at −80°C. Prior to MNase digestion, the cells were resuspended in pre-incubation solution (final 20 mM citric acid, 20 mM Na_2_HPO_4_, 40 mM EDTA, pH 8.0), with aliquots taken for cell-counting. The final volume of resuspension was subsequently adjusted to have 4.5 x 10^7^ cells/ml. The cells were pre-treated with freshly added 2-mercaptoethanol (2-ME, final 30 mM in pre-incubation buffer) for 10 minutes at 30°C, followed by zymolyase treatment in zymolyase buffer (final 1 M sorbitol, 50 mM Tris-HCl (pH 7.5), 10 mM 2-ME and 1 mg/ml 100T zymolyase) for 30-35 minutes [97]. Converted spheroplasts were washed once with cold zymolyase buffer without 2-ME, resuspended in nystatin buffer (final 50 mM NaCl, 1.5 mM CaCl_2_, 20 mM Tris-HCl (pH 8.0), 1 M sorbitol, and 100 ug/ml nystatin (Sigma), and then kept on ice temporarily.

The following MNase digestion was performed for each strain individually. Resuspended spheroplasts were sequentially added into the MNase aliquots (ranged from final 0.0125 to 0.1 U/ml, prepared in nystatin buffer), and incubated at 25°C for 15 minutes. Reactions were stopped by adding 1% SDS/12 mM EDTA (final concentration) [98, 99]. Subsequently, the spheroplasts were treated with RNase (final 0.02 μg/μl) at 37°C for 45 minutes, followed by proteinase K (final 0.4 μg/μl) at 65°C, overnight. The DNA samples were purified with phenol/chloroform extraction, precipitated with ethanol overnight and then resuspended in 1X TE. The samples (2.5 μg) were analyzed with gel electrophoresis (1.2% TAE agarose gel, at 35 V overnight) [97].

## Data availability

The datasets and computer code related to this study are available in the following databases:

- RNA-seq data: Gene Expression Omnibus 163287 (https://www.ncbi.nlm.nih.gov/geo/query/acc.cgi?acc=GSE163287)
- ChIP-seq data: Gene Expression Omnibus 42655 (https://www.ncbi.nlm.nih.gov/geo/query/acc.cgi?acc=GSE42655)

## Acknowledgements

We thank K. Shirahige and Y. Katou for support and technical assistance during collection of the previously published ChIP-Seq data. We thank C. Björkegren for sharing strains and plasmids, and for critical reading of the manuscript.

---

**S1 Fig.**
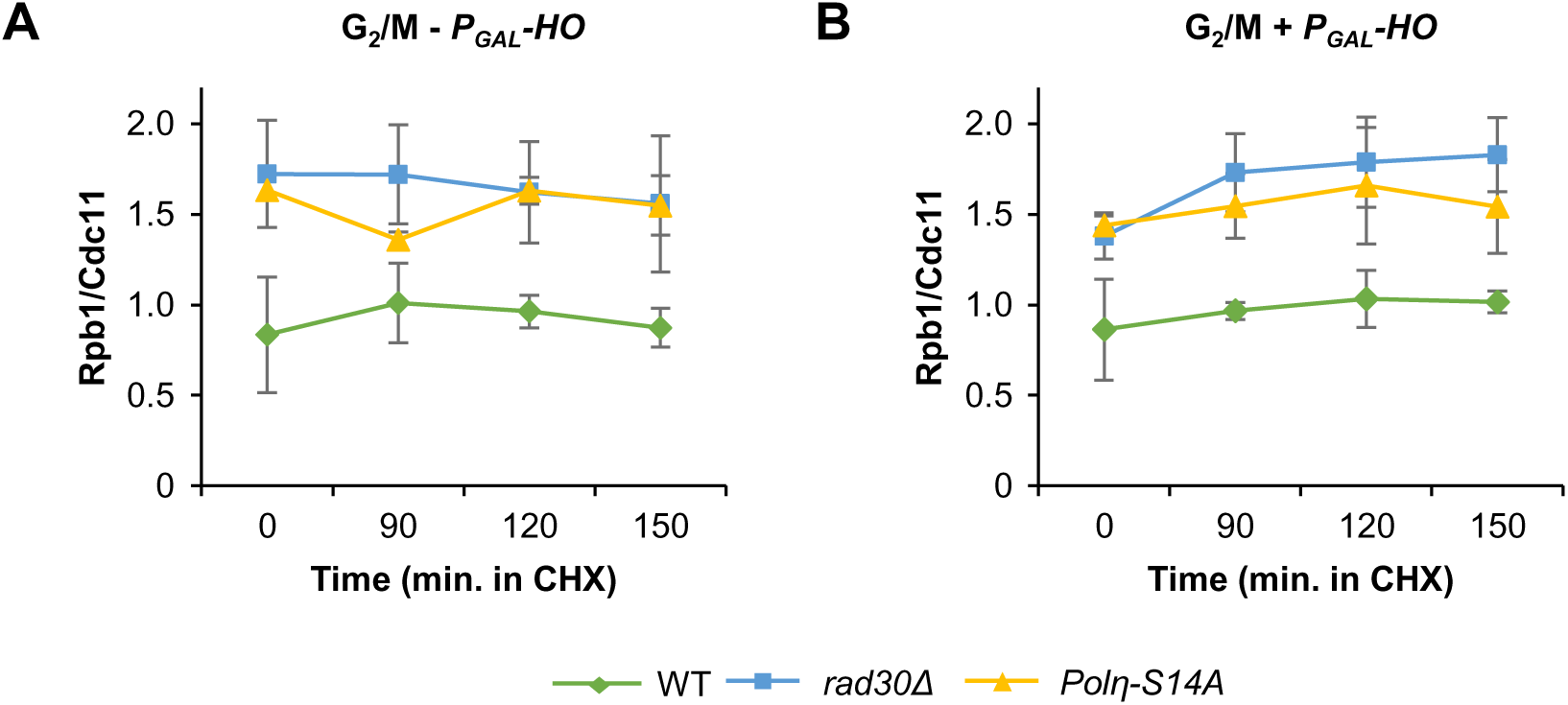
Quantitation of Rpb1 levels. (A-B) Relative amounts of Rpb1 after addition of water (A, control) or galactose (B) to induce *P_GAL_-HO* DSB induction for one-hour, followed by cycloheximide (CHX) chase up to 150 minutes. Western blots from two independent experiments were quantified to compare Rpb1 levels (relative to Cdc11) between the indicated strains.

**S2 Fig.**
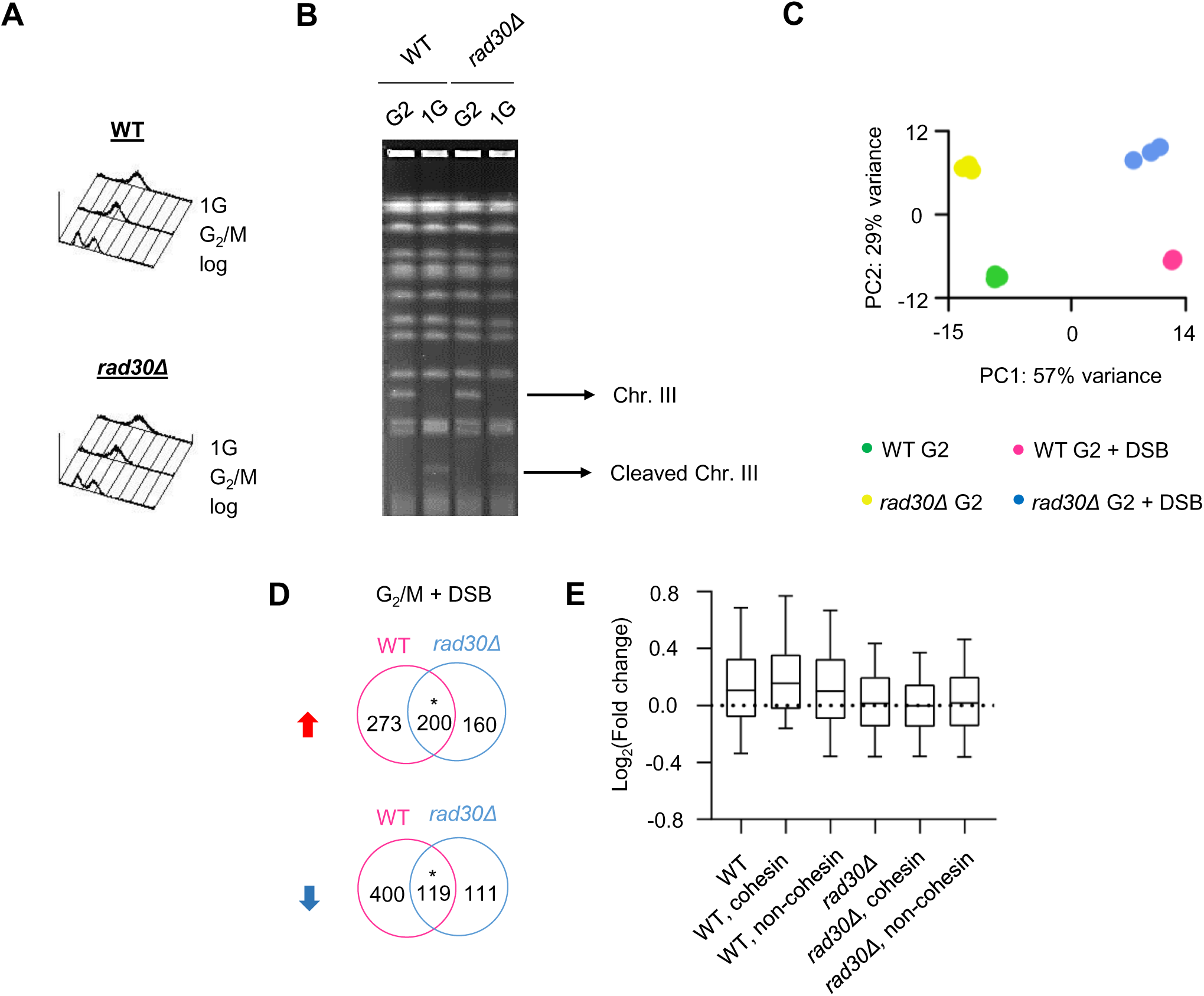
Control experiments for RNA-seq and a differential gene expression analysis in relation to cohesin binding. (A) FACS analysis to confirm benomyl-induced G_2_/M arrest. 1G, 1-hour *GAL*-induction (P*_GAL_-HO*). (B) PFGE analysis to monitor DSB induction on chromosome III. G2, G_2_/M arrest; 1G as in (A). (C) PCA demonstrating distribution of independent data sets between groups and clustering of data sets within groups. (D) Venn diagrams showing overlaps of differentially expressed genes in WT and *rad30Δ* cells after DSB, based on RNA-seq. The red and blue arrows indicate up- and down-regulated genes respectively. Statistical significance of the overlapping genes was evaluated as described in Materials and Methods, with * *p* < 0.001. (E) Expression changes of short genes (< 500 bp) after DSB in WT and *rad30Δ* cells. Genes with or without cohesin enrichment were defined according to the Scc1 ChIP-seq data.

**S3 Fig.**
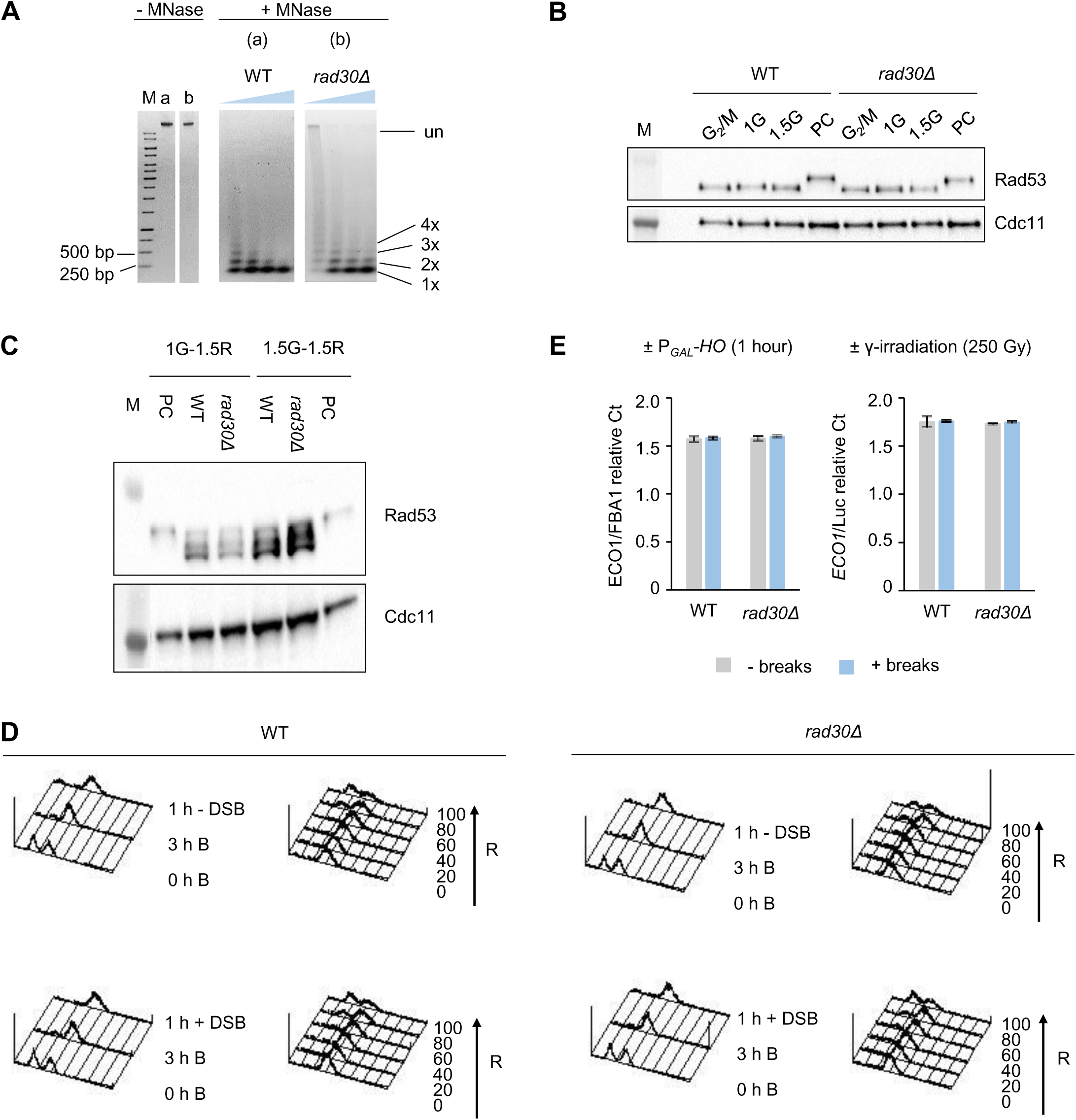
The *rad30Δ* mutant showed increased nucleosome occupancy, but no difference in activation of DNA damage checkpoint and *ECO1* gene expression compared to WT cells. (A) Monitoring nucleosome occupancy based on sensitivity of cells to MNase digestions. The concentrations of MNase were 0, 0.0125, 0.025, 0.05, 0.1 U/ml (final). One representative gel electrophoresis from at least two independent assays performed is shown. The gel images were cropped to show selected samples. M, DNA ladder; Un, undigested; 1x, monomer; 2x, dimer; 3x, trimer; 4x, tetramer. (B) Monitoring activation of the DNA damage checkpoint (phosphorylation of Rad53) after DSB induction with western blot. Galactose was added into the G_2_/M arrested cell cultures to induce P*_GAL_*-*HO* break induction for 1- or 1.5-hour, denoted as 1G or 1.5G. Sample collected from G_2_/M arrested WT cells, treated with phleomycin (final 15 μg/ml) for 1.5 hours was included as positive control (PC). Cdc11 was used as loading control. M, protein marker. (C) Monitoring activation of DNA damage checkpoint during DSB recovery. DSB was induced for 1- or 1.5-hour, as in (B). The cells were then allowed to recover in YEP media supplemented with glucose and benomyl for another 1.5 hour (1.5 R) at 35°C, to mimic the damage-induced cohesion assay. 1G, 1.5G, PC, M as in (B). Cdc11 was used as loading control. (D) FACS analyses of cell cycle progression in WT and *rad30Δ* cells, at indicated time points after release into YEP media supplemented with glucose to recover from DSB induction. Samples without DSBs were included as control. B, benomyl; R, recovery. (E) *ECO1* gene expression in G_2_/M arrested WT and *rad30Δ* cells ± P*_GAL_-HO* (left) and ± γ-irradiation (right). The relative gene expression was measured by RT-qPCR. *FBA1* was used as a reference gene for the ± P*_GAL_-HO* samples. Purified total RNA (0.65 μg) was spiked in with 1 ng luciferase control RNA (Promega) prior to cDNA synthesis for the ± γ-irradiation samples. Error bars indicate the mean ± STDEV of at least two independent experiments.

**S4 Fig.**
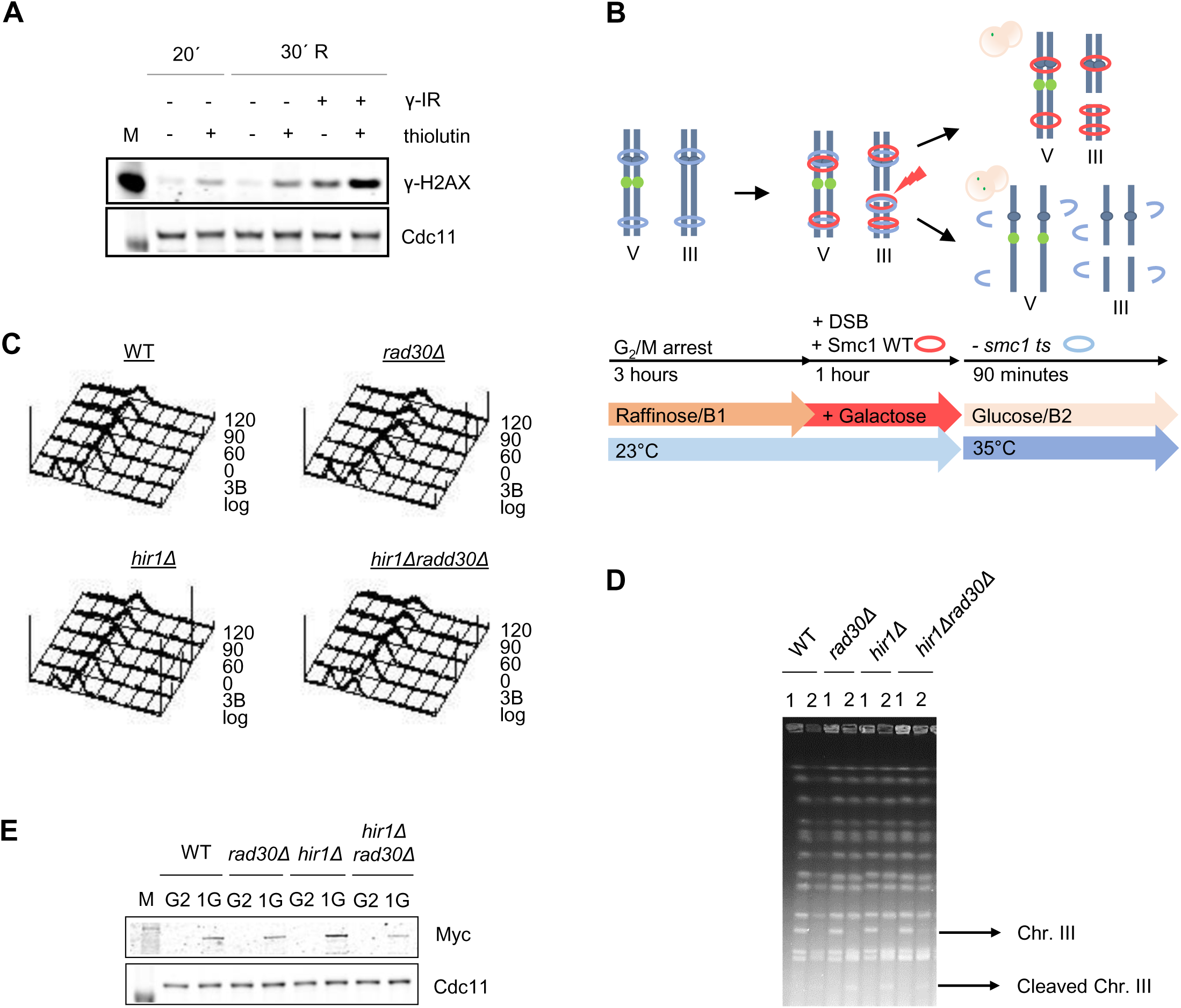
The method and related control experiments for a typical damage-induced cohesion assay. (A) Detection of early DNA damage response (γ-H2AX) in thiolutin treated cells. G_2_/M arrested WT cells were treated with 20 µg/ml thiolutin (final) for 20 minutes, denoted as 20’. The culture was then split for ± γ-irradiation, and allowed to recover for 30 minutes in the presence of thiolutin after ± γ-irradiation (30’ R). Gamma-irradiated cells without thiolutin treatment were included as control. Cdc11 was used as loading control. M, protein marker. (B) Damage-induced cohesion assay performed with *GAL* induced DSBs on chromosome III (P*_GAL_*-*HO*). Strains harboring the temperature sensitive *smc1-259* allele are arrested in G_2_/M by addition of benomyl (B). Galactose is then added for expression of ectopic P*_GAL_*-*SMC1*-*MYC* (Smc1 WT) and induction of DSBs, for 1-hour. The temperature is then raised to 35°C, restrictive to the *smc1-259* allele, for disruption of S-phase cohesion (blue rings). The Tet-O/TetR-GFP system (green dots) is used to monitor damage-induced cohesion (red rings) on chr. V. Chr., chromosome; III, three; V, five. B1 and 2 indicate replacement of media with freshly prepared benomyl. (C) FACS analysis to confirm G_2_/M arrest during the time course of a typical damage-induced cohesion assay. 3B, 3-hour benomyl arrest. (D) PFGE analysis to detect DSB induction on chromosome III. 1, G_2_/M arrest; 2, 1-hour *GAL*-induction (P*_GAL_-HO* and P*_GAL_-SMC1-MYC*). (E) Western blot to check expression of the *GAL* promoter driven ectopic Smc1-Myc protein. G2, G_2_/M arrest; 1G, 1-hour *GAL*-induction as in (D). M and Cdc11 as in (A).

**Fig S5.**
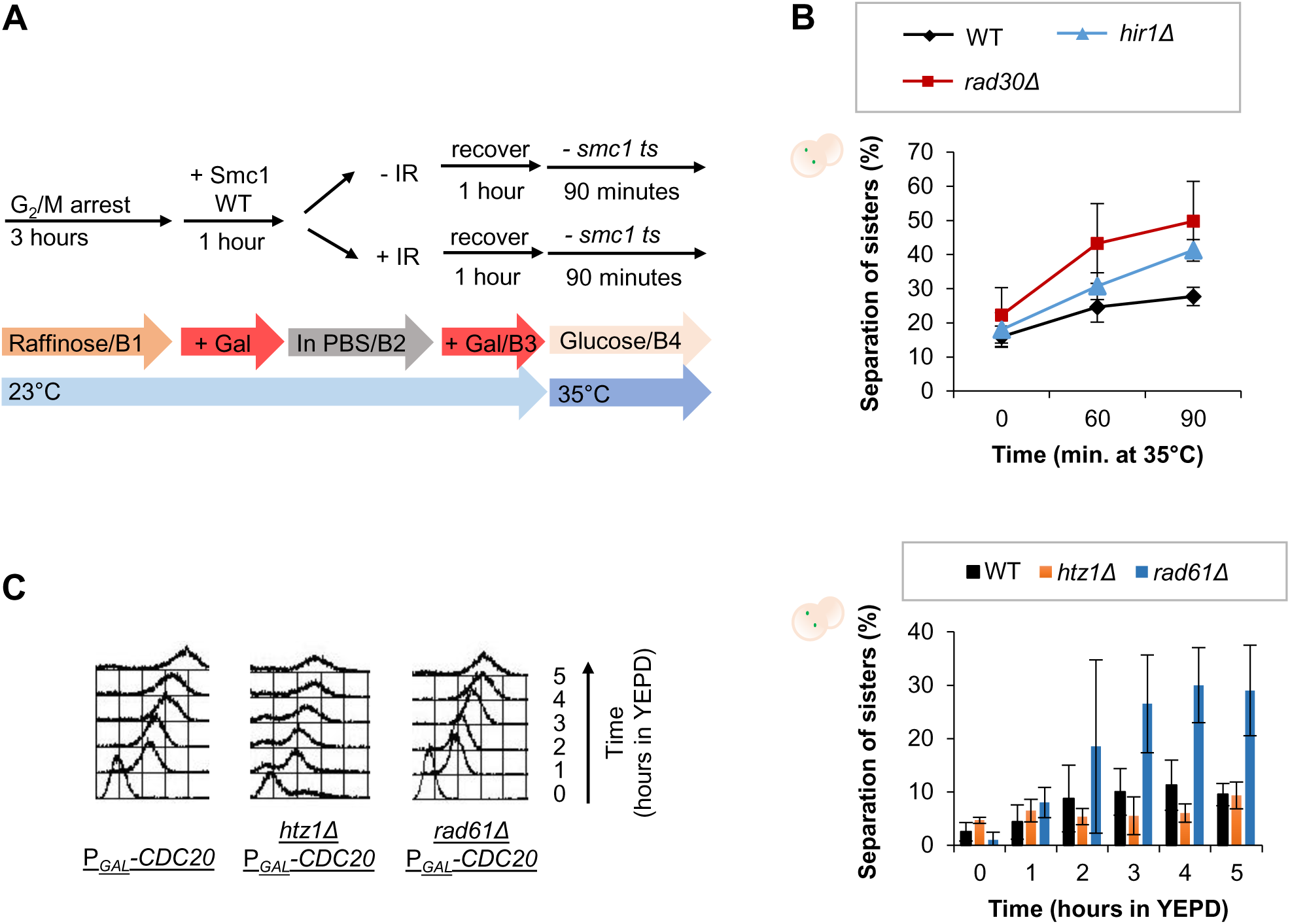
Damage-induced cohesion assay performed with γ-irradiation and the maintenance of sister chromatid cohesion in *htz1Δ* cells. (A) Damage-induced cohesion assay performed with γ-irradiation. Formation of damage-induced cohesion is monitored on chr. V with the same Tet-O/TetR-GFP system, as in S4B Fig, with slight differences in the experimental procedure. Strains with *smc1-259* background are arrested in G_2_/M by addition of benomyl (B), expression of ectopic P*_GAL_*-*SMC1*-*MYC* (Smc1 WT) is then induced by addition of galactose. The cells are subsequently pelleted, resuspended in 1X PBS supplemented with benomyl. The resuspension is split in one half for irradiation, and half as non-irradiated control. After irradiation, both ± irradiated cells are recovering in YEP media supplemented with galactose and benomyl. Subsequently, the media is changed to YEP containing glucose and benomyl, and the temperature raised to 35°C, to monitor formation of damage-induced cohesion. (B) Damage-induced cohesion assay of the *hir1Δ* mutant in response to γ-irradiation, performed as depicted in (A). Means ± STDEV from at least two independent experiments are shown. (C) Sister chromatid cohesion maintenance of the *htz1Δ* mutant under prolonged G_2_/M arrest. The cells were initially synchronized in G_1_ by α-factor in YEP media containing galactose. Expression of P*_GAL_-CDC20* was then shut off by switching the carbon source to glucose (YEPD), which resulted in the subsequent prolonged G_2_/M arrest as monitored by FACS (left panel). Sister chromatid separation was monitored at the *URA3* locus on Chr. V by the TetO/TetR-GFP system. Means ± STDEV from at least two independent experiments are shown (right panel). A *rad61Δ* mutant with known high sister separation under prolonged G_2_/M arrest was included as control. Means ± STDEV from at least two independent experiments are shown. Parts of the results from the same experiments were previously published [28]. Chr., chromosome.

**Fig S6.**
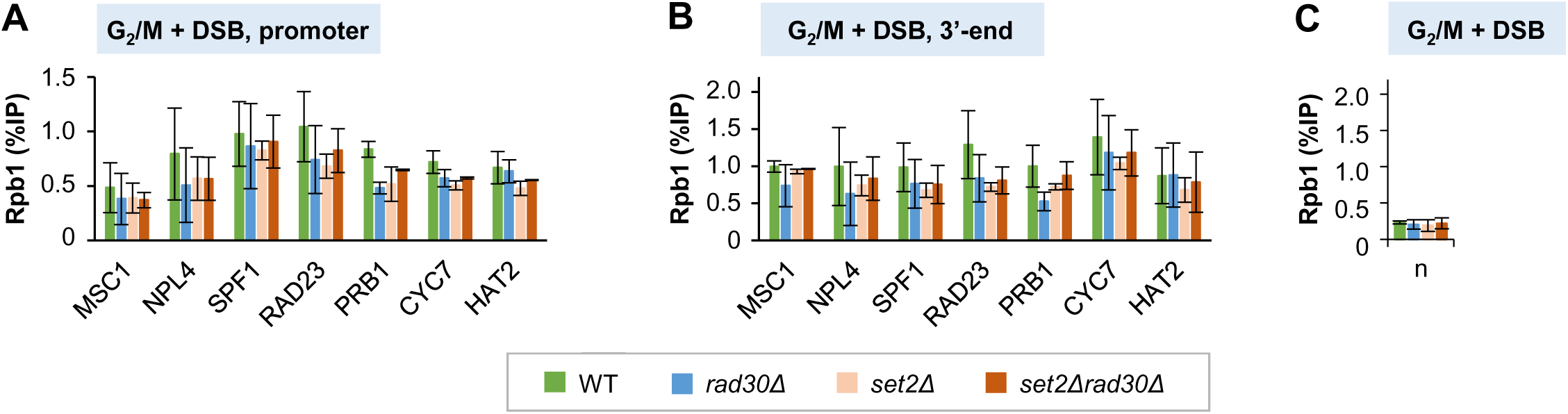
Chromatin association of Rpb1 in the *set2Δrad30Δ* double mutant after DSB induction. (A-C) ChIP-qPCR analyses to determine chromatin association of Rpb1 at promoters and 3’-ends of selected genes, in G_2_/M arrested cells after DSB induction. The same genes as in Fig 7C-7E were analyzed. Error bars indicate the mean ± STDEV of at least two independent experiments. n, low-binding control (n2 in Fig 1B).

**Fig S7.**
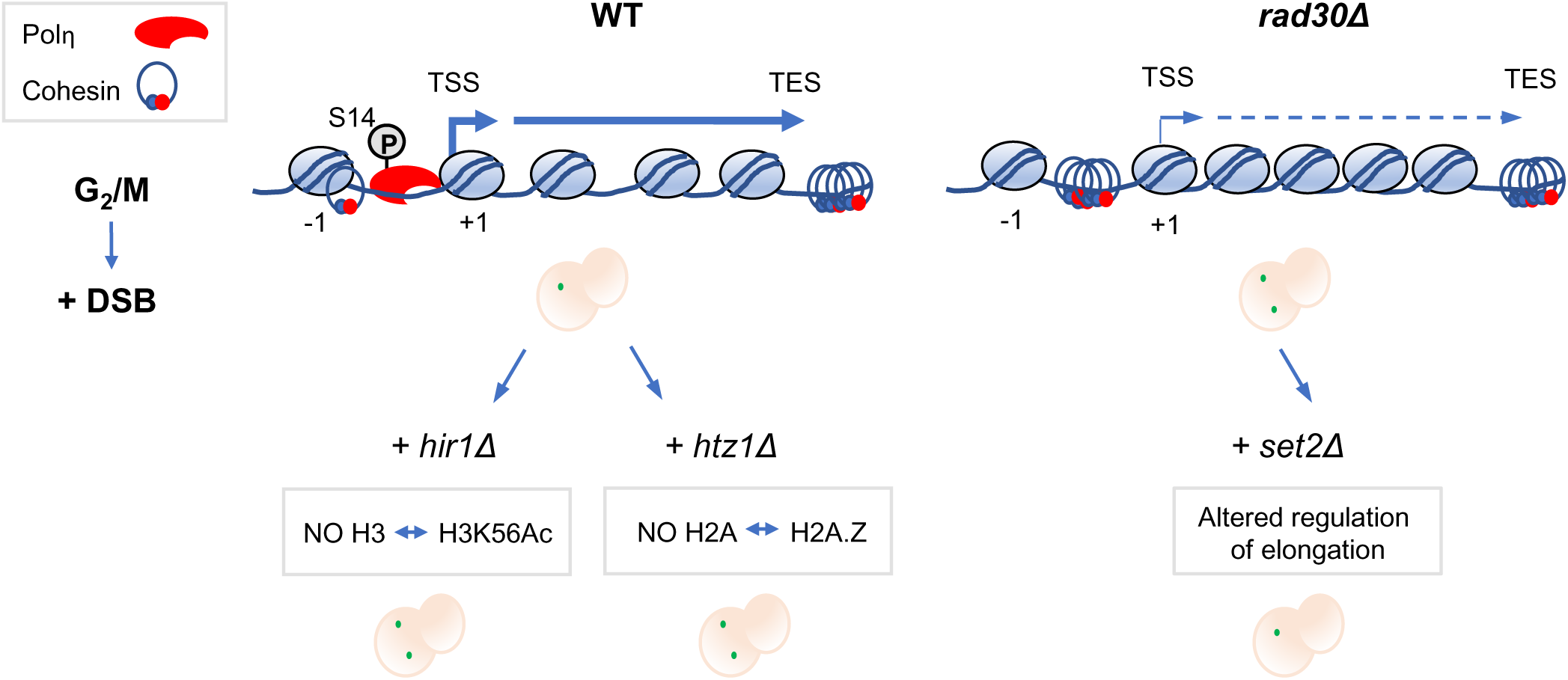
A summary of the main results and a proposed model. In G_2_/M arrested WT cells, genes belonging to the positive transcription regulation and chromatin assembly pathways are enriched compared to *rad30Δ* cells. Reduced chromatin assembly in *rad30Δ* cells results in less dynamic chromatin, indicated by additional nucleosomes. Deregulated transcription and sensitivity to elongation inhibitors in *rad30Δ* cells are indicated by thin arrows over the TSS and ORF. Histone exchange between H3 and H3K56Ac at promoter regions is reduced in the *hir1Δ* mutant, while histone exchange of H2A for H2A.Z at the +1 nucleosome is prevented in the *htz1Δ* mutant, hampering transcriptional regulation. Both mutants were deficient in damage-induced cohesion. In contrast, deletion of *SET2* compensated for reduced transcriptional capacity of the *rad30Δ* mutant, and suppressed the lack of damage-induced cohesion in *rad30Δ* cells. Taken together, histone exchange during transcription is proposed to facilitate formation of damage-induced cohesion. This process is perturbed in *rad30Δ* cells, which functionally associates with transcription, thereby negatively affecting generation of damage-induced cohesion. Cells with a single green dot indicates established damage-induced cohesion while cells with two dots indicates lack of damage-induced cohesion. ORF, open reading frame.

**S1 Table.**
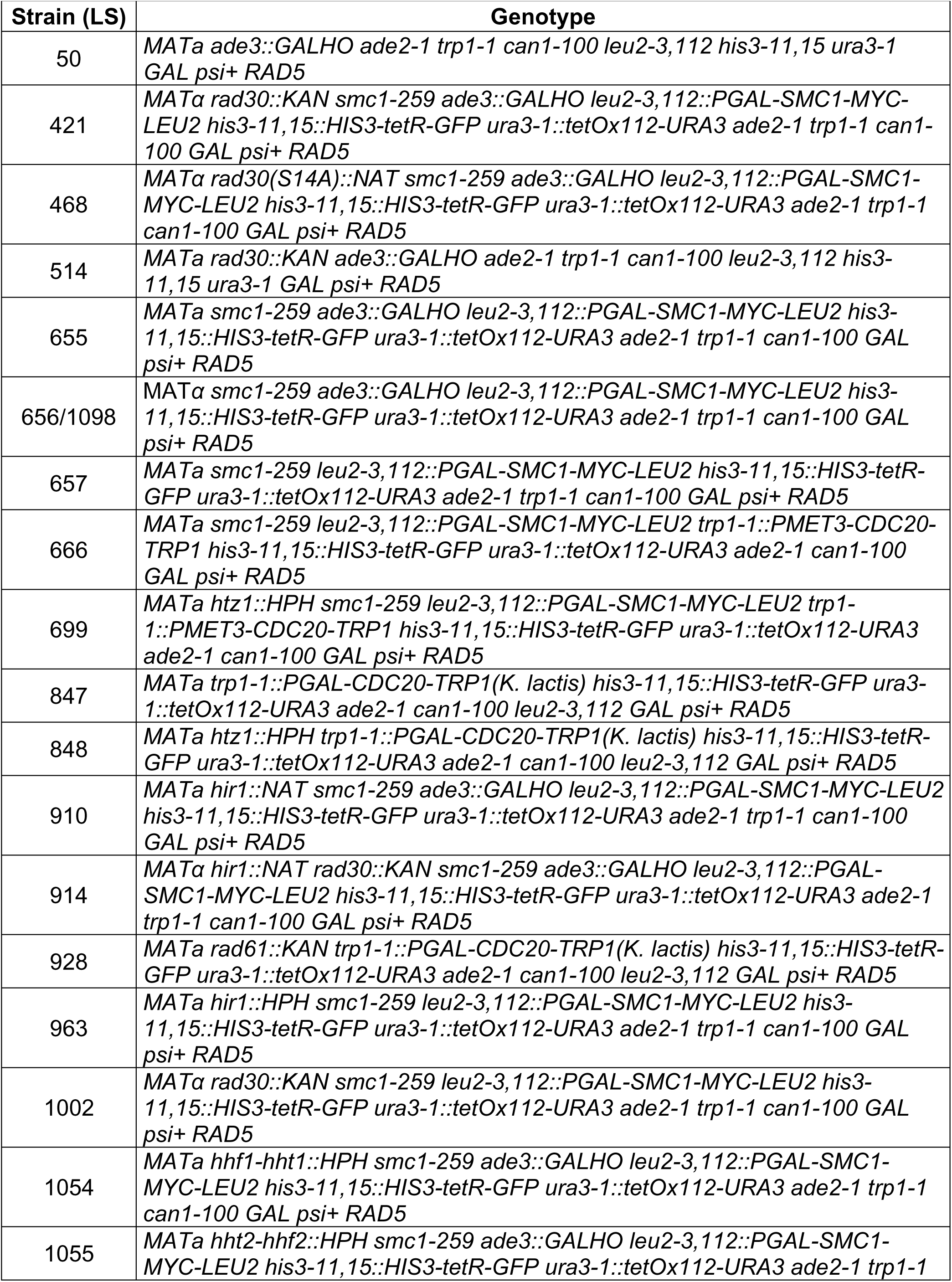
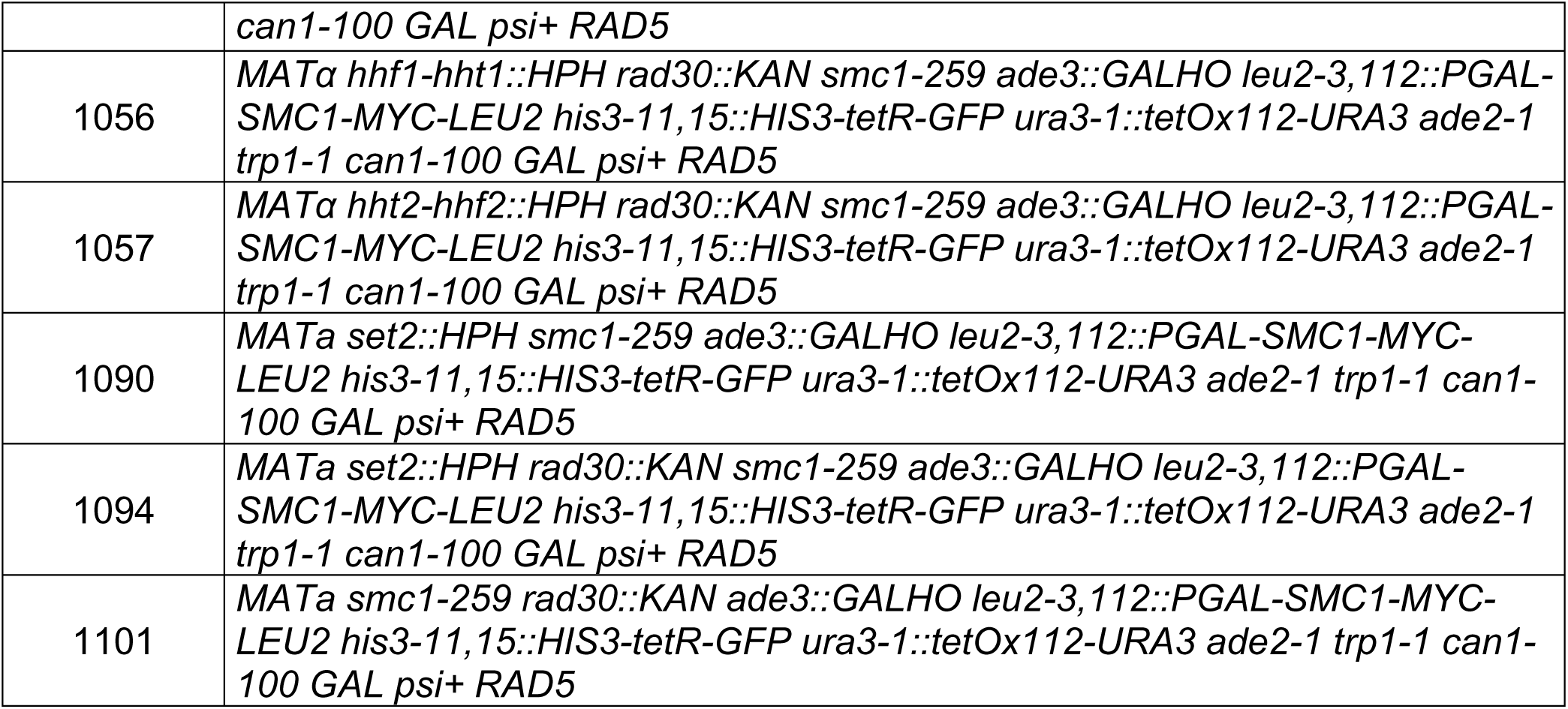
Strains used in this study.

**S2 Table.** Information on primary antibodies used.

| **Antibody** | **Company** | **Catalog #** |
| --- | --- | --- |
| anti-Rpb1 (8WG16) | Santa Cruz Biotechnology | sc-56767 |
| anti-Cdc11 (y-415) | Santa Cruz Biotechnology | sc-7170 |
| anti-Rad53 | Abcam | ab104232 |
| anti-c-myc | Roche | 11667203001 |
| anti-Htz1 | Active Motif | 39647 |
| anti-Histone H2A (phospho S129) | Abcam | ab15083 |

**S3 Table.**
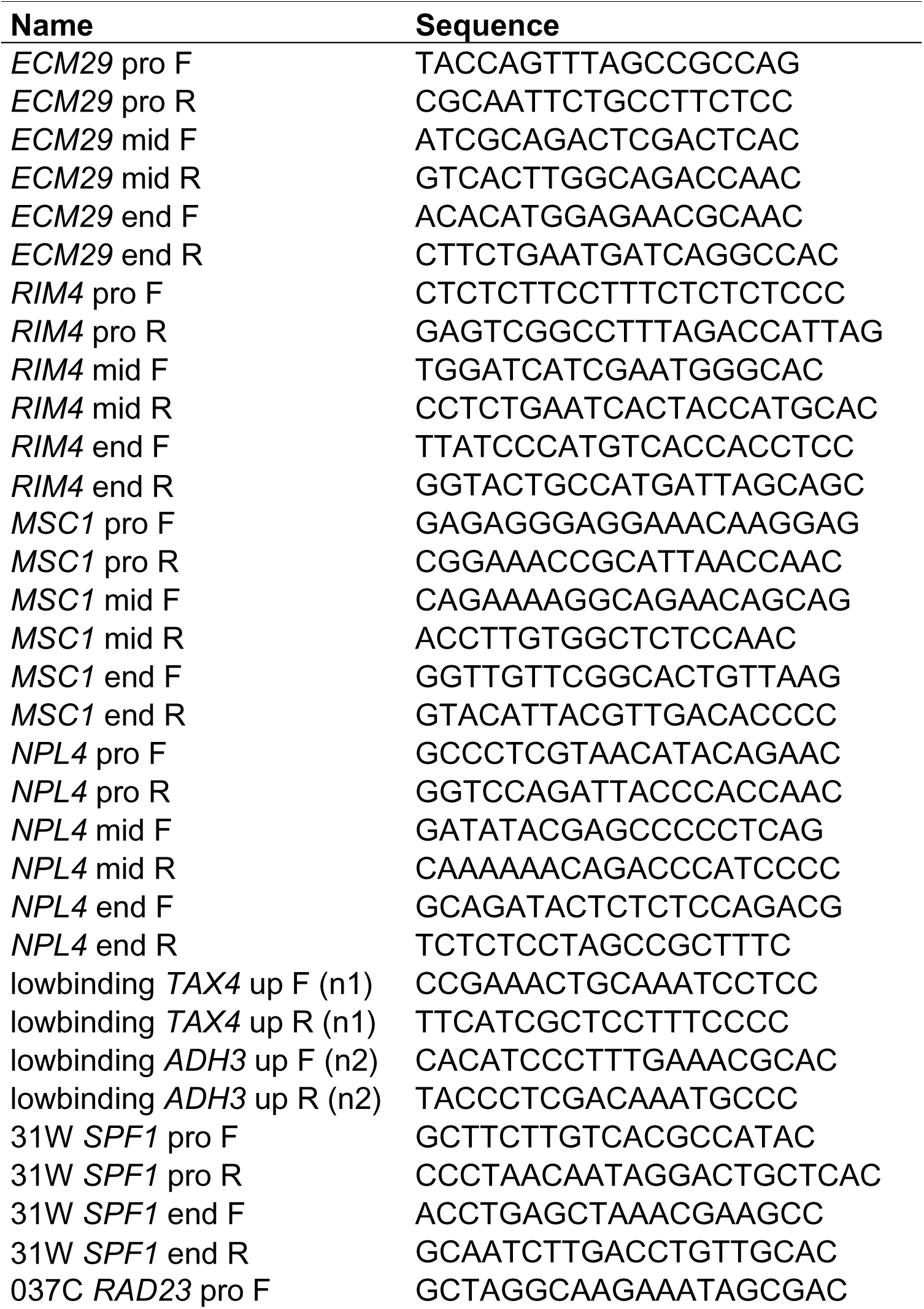
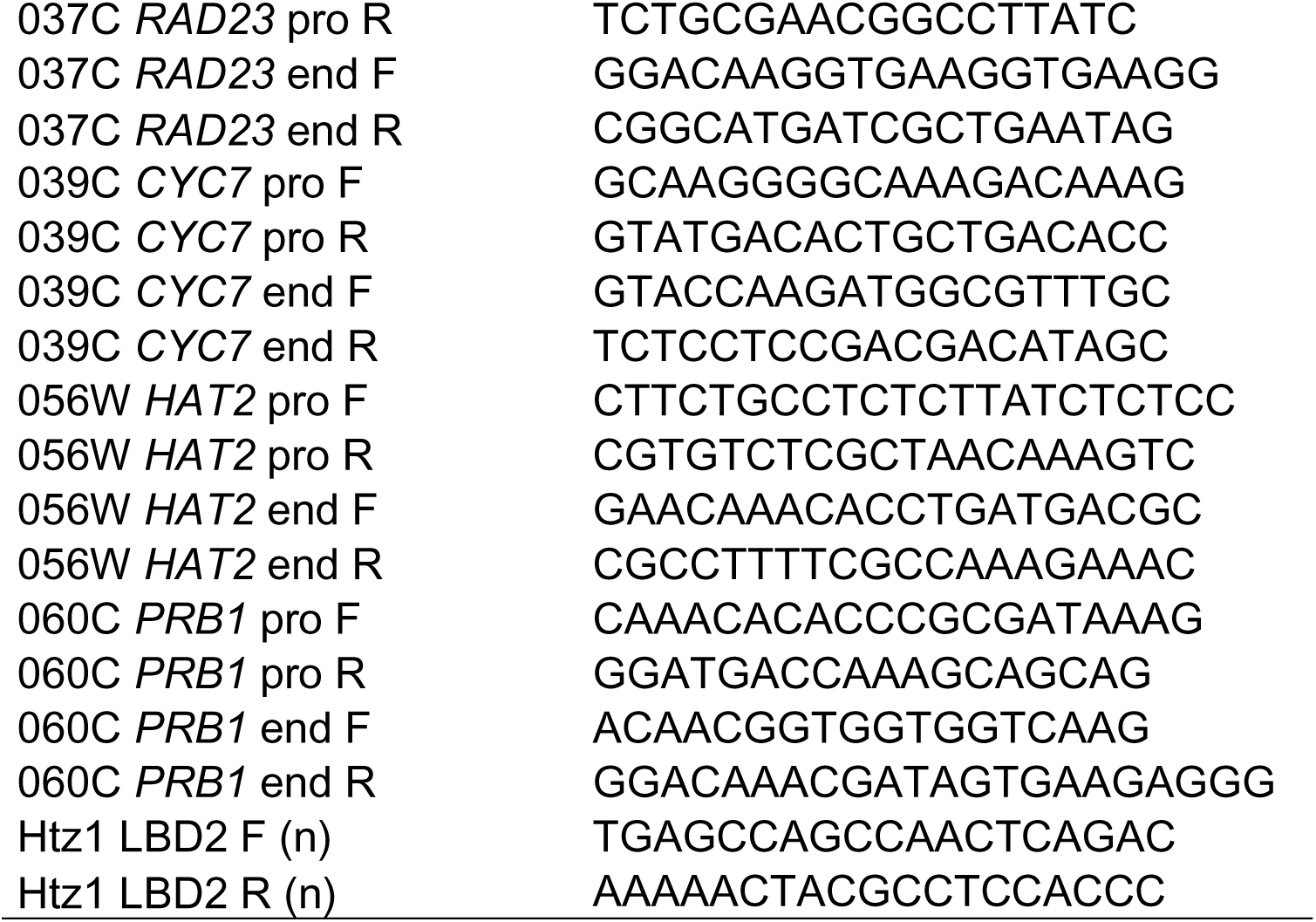
Primers used in ChIP-qPCR.

